# Quantitative profiling of native RNA modifications and their dynamics using nanopore sequencing

**DOI:** 10.1101/2020.07.06.189969

**Authors:** Oguzhan Begik, Morghan C Lucas, Leszek P Pryszcz, Jose Miguel Ramirez, Rebeca Medina, Ivan Milenkovic, Sonia Cruciani, Huanle Liu, Helaine Graziele Santos Vieira, Aldema Sas-Chen, John S Mattick, Schraga Schwartz, Eva Maria Novoa

## Abstract

A broad diversity of modifications decorate RNA molecules. Originally conceived as static components, evidence is accumulating that some RNA modifications may be dynamic, contributing to cellular responses to external signals and environmental circumstances. A major difficulty in studying these modifications, however, is the need of tailored protocols to map each modification type individually. Here, we present a new approach that uses direct RNA nanopore sequencing to identify and quantify RNA modifications present in native RNA molecules. First, we show that each RNA modification type results in a distinct and characteristic base-calling ‘error’ signature, which we validate using a battery of genetic strains lacking either pseudouridine (Y) or 2’-O-methylation (Nm) modifications. We then demonstrate the value of these signatures for *de novo* prediction of Y modifications transcriptome-wide, confirming known Y-modified sites as well as uncovering novel Y sites in mRNAs, ncRNAs and rRNAs, including a previously unreported Pus4-dependent Y modification in yeast mitochondrial rRNA, which we validate using orthogonal methods. To explore the dynamics of pseudouridylation across environmental stresses, we treat the cells with oxidative, cold and heat stresses, finding that yeast ribosomal rRNA modifications do not change upon environmental exposures, contrary to the general belief. By contrast, our method reveals many novel heat-sensitive Y-modified sites in snRNAs, snoRNAs and mRNAs, in addition to recovering previously reported sites. Finally, we develop a novel software, *nanoRMS*, which we show can estimate per-site modification stoichiometries from individual RNA molecules by identifying the reads with altered current intensity and trace profiles, and quantify the RNA modification stoichiometry changes between two conditions. Our work demonstrates that Y RNA modifications can be predicted *de novo* and in a quantitative manner using native RNA nanopore sequencing.

## INTRODUCTION

RNA modifications are chemical moieties that decorate RNA molecules, expanding their lexicon. By coupling antibody immunoprecipitation or chemical probing with next-generation sequencing (NGS), transcriptome-wide maps of several RNA modifications have been constructed, including N6-methyladenosine (m^6^A) ^1,2^, pseudouridine (Y) ^3–6^, 5-methylcytosine (m^5^C) ^7,8^, 5-hydroxymethylcytosine (hm^5^C) ^9^, 1-methyladenosine (m^1^A) ^10,11^, N3-methylcytosine (m^3^C) ^12^, N4-acetylcytosine (ac^4^C) ^13,14^ and 7-methylguanosine (m^7^G) ^15,16^. These studies have revealed that RNA modifications play a pivotal role in a large variety of cellular processes, including regulation of cellular fate ^17^, sex determination ^18^ and cellular differentiation ^19^, among others.

Despite these advances, a fundamental challenge in the field is the lack of a generic approach for mapping diverse RNA modification types simultaneously ^20–23^. Currently, customized protocols must be individually set up and optimized for each RNA modification type, leading to experimental designs in which the RNA modification type to be studied is chosen beforehand, hindering the ability to characterize the plasticity of the epitranscriptome in a systematic and unbiased manner in response to different conditions. Moreover, even in those cases where a selective antibody or chemical is available, NGS-based methods are often not quantitative (i.e. cannot solve the ‘stoichiometry’ problem), have high false positive rates ^21^, are inconsistent when using distinct antibodies ^24^, are unable to produce maps for highly repetitive regions, cannot provide information regarding the co-occurrence of distant modifications in same transcripts, do not provide isoform-specific information, and require multiple ligations steps and extensive PCR amplification during the library preparation, introducing undesired biases in the sequencing data ^25^.

A promising alternative to NGS-based technologies that can, in principle, overcome these limitations is the direct RNA sequencing platform developed by Oxford Nanopore Technologies (ONT), which has the potential to detect virtually any given RNA modification present in native RNA molecules ^20,26,27^. Algorithms to detect RNA modifications have been made available in the last few months ^28–30^, many of which rely on the use of systematic base-calling ‘errors’ caused by the presence of RNA modifications. However, to date the vast majority of efforts have been devoted to the detection of m^6^A modifications ^29–33^, and it is largely unknown whether other modifications of RNA bases may be distinguishable from their unmodified counterparts using this technology. Thus, a systematic, multiplexed and unbiased approach that can map and quantify diverse RNA modifications simultaneously in full-length molecules is currently missing.

Here, we examine the *S. cerevisiae* coding and non-coding transcriptome at single molecule resolution using native RNA nanopore sequencing. We find that most RNA modifications are characterized by systematic base-calling errors, and that the signature of these base-calling ‘errors’ can be used to identify the underlying RNA modification type. For example, we find that pseudouridine typically appears in the form of U-to-C mismatches, whereas m^5^C modifications appear in the form of insertions. We then exploit the identified signatures to *de novo* predict RNA modifications in rRNAs, finding two previously unreported Y modifications in mitochondrial rRNA, which we confirm using CMC-probing coupled to nanopore sequencing (nanoCMC-seq). We demonstrate that one of these novel Y modifications (15s:Y854) is placed by the enzyme Pus4, which was previously thought to pseudouridylate only mRNAs and tRNAs ^4^. Moreover, we show that once the Y RNA modifications have been accurately predicted using base-calling ‘errors’, the stoichiometry of a given Y- or Nm-modified site can be estimated by clustering per-read features (current intensities and trace) of the modified regions.

We then explore the dynamics of RNA modifications present in non-coding RNAs. It has been proposed that differential rRNA modifications may constitute a source of ribosomal heterogeneity ^34–36^, leading to fine tuning of the ribosomal function and ultimately proteome output. Indeed, previous studies have shown that temperature changes affect rRNA pseudouridylation levels at specific sites, suggesting that cells may be able to generate compositionally distinct ribosomes in response to environmental cues ^4,37,38^. Similarly, alterations in the stoichiometry of 2’-O-methylation (Am, Cm, Gm, Um) ^39–41^ and pseudouridylation (Y) ^34–36^ have been shown to affect translation initiation of mRNAs containing internal ribosome entry sites (IRES) ^42,43^. Here we re-examine this question using direct RNA sequencing, and characterize the RNA modification dynamics in rRNAs, snRNAs and snoRNAs upon a battery of environmental cues, translational repertoires and genetic strains. Contrary to expectations, we find that none of the environmental stresses tested lead to significant changes in the ribosomal epitranscriptome. By contrast, our method does recapitulate previously reported heat-dependent Y snRNA modifications, as well as identifies novel heat-sensitive sites in snRNAs and snoRNAs.

Finally, we develop a novel algorithm, *nanoRMS*, which we demonstrate can predict Y RNA modifications *de novo*, as well as estimate the stoichiometry of modification both in highly-modified and lowly-modified Y and Nm sites, and illustrate its applicability *in vivo* across diverse types of RNA molecules, including rRNAs, sn/snoRNAs and mRNAs. To this end, we first systematically examine how the choice of distinct per-read features (signal intensity, dwell time and trace) affects our ability to accurately predict RNA modification stoichiometry from individual read information. Secondly, we benchmark how the machine learning algorithm choice affects the performance of the predictions. Thirdly, we assess the robustness of the distinct algorithms and feature combinations upon diverse ranges of RNA modification stoichiometries. Fourthly, we demonstrate its applicability *in vivo*, by showing that it can be applied to highly-modified non-coding RNA molecules such as rRNAs, snRNAs and snoRNAs, as well as to lowly-modified mRNA molecules. Our approach recapitulates known Pus1-dependent, Pus4-dependent and heat stress-dependent mRNA sites, as well as reveals novel Y mRNA sites that had not been previously reported. Altogether, our work establishes a framework for the study of RNA modification dynamics using direct RNA sequencing, opening novel avenues to study the plasticity of the epitranscriptome at single molecule resolution.

## RESULTS

### Detection of RNA modifications in direct RNA sequencing data is strongly dependent on base-calling and mapping algorithms

Previous studies have shown that N6-methyladenosine (m^6^A) RNA modifications can be detected in the form of non-random base-calling ‘errors’ in direct RNA sequencing datasets ^29–33^. However, it is unclear how these ‘errors’ may vary with the choice of base-calling and mapping algorithms, and consequently, affect the ability to detect and identify RNA modifications. To systematically determine the accuracy of commonly used algorithms for direct RNA base-calling, as well as to assess their ability to detect RNA modifications in the form of base-calling ‘errors’ ^29^, we compared their performance on *in vitro* transcribed RNA sequences which contained all possible combinations of 5-mers, referred to as ‘curlcakes’ (CCs) ^29^, that included: (i) unmodified nucleosides (UNM), (ii) N6-methyladenosine (m^6^A), (iii) pseudouridine (Y), (iv) N5-methylcytosine (m^5^C), and (v) N5-hydroxymethylcytosine (hm^5^C) (**Figure 1A**). In addition, a sixth dataset containing unmodified short RNAs (UNM-S), with median length of 200 nucleotides, was included in the analysis to assess the effect of input sequence length in base-calling (see *Methods*). Each dataset was base-called with two distinct algorithms (*Albacore* and *Guppy*), and using two different versions for each of them, namely: (i) *Albacore* version 2.1.7 (AL 2.1.7); (ii) its latest version, *Albacore* 2.3.4 (AL 2.3.4); (iii) *Guppy* 2.3.1 (GU 2.3.1); and (iv) a more recent version of the latter base-caller, *Guppy* 3.0.3 (GU 3.0.3), which employs a flip-flop algorithm. We found that the latest version of *Albacore* (2.3.4) base-called 100% of sequenced reads in all 6 datasets, whereas its previous version did not (average of 90.8%) (**Figure 1B**). In contrast, both versions of *Guppy* (2.3.1 and 3.0.3) produced similar results in terms of percentage of base-called reads (98.71% and 98.75%, respectively) (**Table S1**).

**Figure 1.**
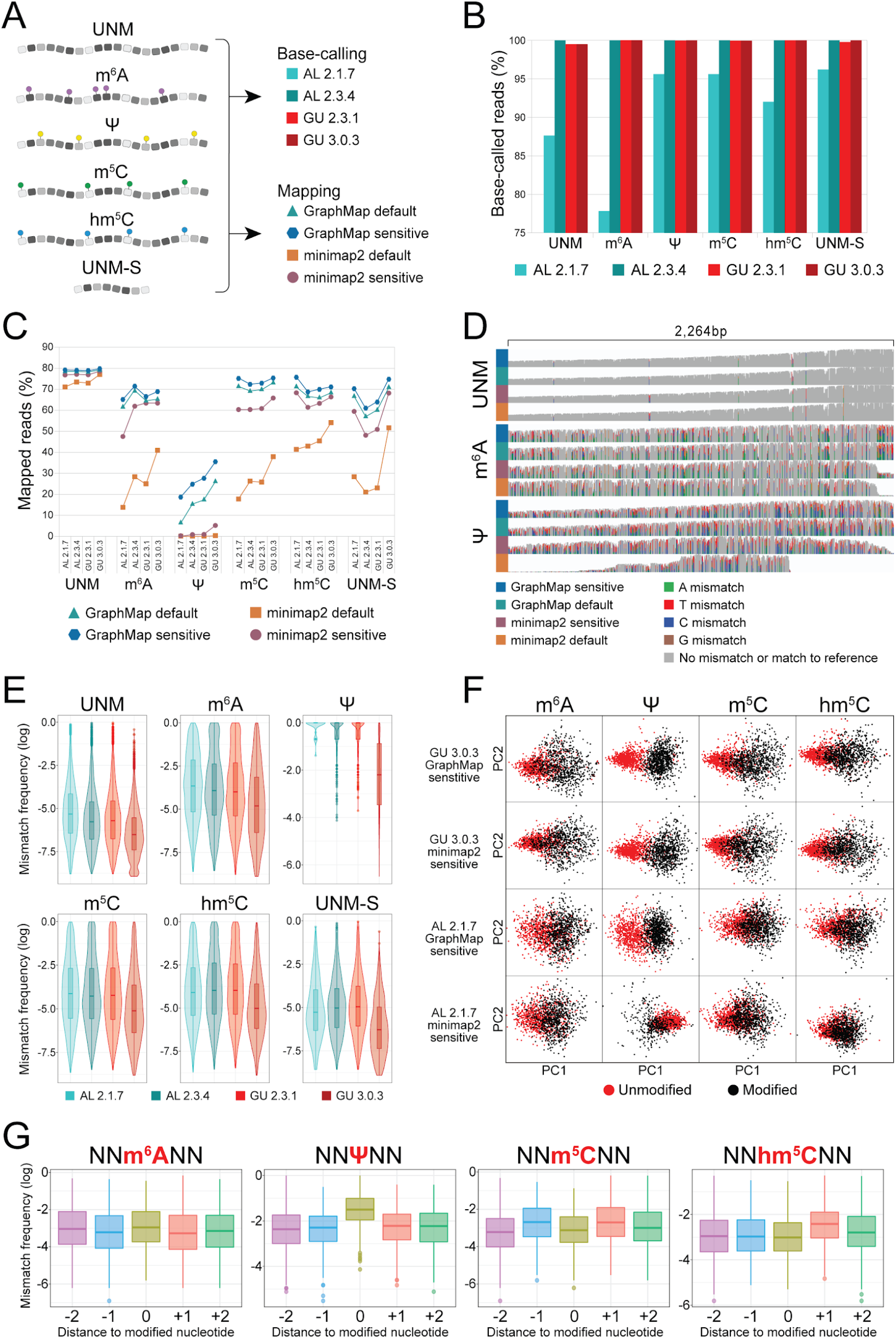
Systematic analysis of base-calling and mapping algorithms for the detection of RNA modifications in direct RNA sequencing datasets. **(A)** Overview of the synthetic constructs used to benchmark the algorithms, which included both unmodified (UNM and UNM-S) and modified (m^6^A, m^5^C, hm^5^C and Y) sequences. For each dataset, we performed: i) comparison of base-calling algorithms, ii) comparison of mapping algorithms, iii) detection of RNA modifications using base-called features and iv) comparative analysis of features to distinguish similar RNA modifications. **(B)** Barplots comparing the percentage of base-called reads using 4 different base-calling algorithms in 6 different unmodified and modified datasets. **(C)** Relative proportion of base-called and mapped reads using all possible combinations (16) of base-callers and mappers included in this study, for each of the 6 datasets analyzed. **(D)** IGV snapshots illustrating the differences in mapping for 3 distinct datasets: UNM, m^6^A-modified and Y-modified when base-called with GU 3.0.3. Mismatch frequencies greater than 0.1 have been colored, grey represents match to reference. **(E)** Comparison of global mismatch frequencies using different base-calling algorithms, for the 6 datasets analyzed. Box, first to last quartiles; whiskers, 1.5x interquartile range; center line, median; points, outliers; violin, distribution of density. **(F)** Principal Component Analysis (PCA) using as input the base-calling error features of quality, mismatch frequency and deletion frequency in positions −2, −1, 0, 1 and 2, for all datasets base-called with GU 3.0.3 and AL 2.1.7 and mapped with GraphMap and minimap2 on sensitive settings. Only k-mers that contained a modification at position 0 were included in the analysis, and the equivalent set of unmodified k-mers was used as a control. **(G)** Mismatch frequency of each position of the 5-mers centered in the modified position (position 0). Box, first to last quartiles; whiskers, 1.5x interquartile range; center line, median; points, outliers. See also Figure S1.

We then assessed whether the choice of mapper might affect the ability to detect RNA modifications. To this end, we employed two commonly used long-read mappers, *minimap2* ^44^ and *GraphMap* ^45^, using either ‘default’ or ‘sensitive’ parameter settings (see *Methods*). Strikingly, we found that the choice of mapper, as well as the parameters used, severely affected the final number of mapped reads for each dataset (**Figure 1C,** see also **Table S1**). The most extreme case was observed with the Y-modified dataset, where *minimap2* was unable to map the majority of the reads (0-0.3% mapped reads) (**Figure 1C,D,** see also **Figure S1A**). By contrast, *GraphMap* ‘sensitive’ was able to map 35.5% of Y-modified base-called reads, proving to be a more appropriate choice for highly modified datasets. To ascertain whether an increase in the number of base-called and mapped reads was at the expense of decreased accuracy, we assessed the sequence identity percentage (as a read-out of accuracy), finding that *GraphMap* outperforms *minimap2* with only a minor loss in accuracy (3%) (**Figure S1B,** see also **Table S2**).

### Base-calling ‘error’ signatures can be used to predict RNA modification type

While base-calling ‘errors’ can be used to identify m^6^A RNA modified sites ^29,30,32^, whether this approach is applicable for the detection of other RNA modifications, and whether these signatures could be employed to distinguish among distinct RNA modification types, is largely unknown. To this end, we systematically characterized the base-calling errors caused by the presence of m^6^A, Y, m^5^C and hm^5^C. We found that, regardless of the base-caller and mapper settings used, modified RNA sequences presented decreased quality scores (**Figure S1C-E**) and higher mismatch frequencies (**Figure 1E**), being these differences more prominent in Y-modified datasets. Principal component analysis of base-calling ‘errors’ of each modified dataset (m^6^A, Y, m^5^C and hm^5^C) -relative to unmodified- showed that this difference was greatest in Y-modified datasets (**Figure 1F**), and maximized in datasets that were base-called with GU 3.0.3. Thus, we find that all four RNA modifications can be detected in direct RNA sequencing data; however, their detection is severely affected by the choice of both base-calling and mapping algorithms, and varies depending on the RNA modification type.

We then examined whether the base-called ‘errors’ observed in modified and unmodified datasets occured in the modified position. We found that both m^6^A and Y modifications led to increased mismatch frequencies at the modified site (**Figure 1G**), mainly in the form of U-to-C mismatches in the case of Y modifications (**Figure S1F**). By contrast, m^5^C and hm^5^C modifications did not appear in the form of increased mismatch frequencies at the modified site; rather, these modifications appeared in the form of increased mismatch frequencies in the neighboring residues (position −1 and +1 in the case of m^5^C modifications; position +1 in hm^5^C) (**Figure 1G**). Moreover, the observed base-called ‘error’ signatures of m^5^C and hm^5^C were also dependent on the sequence context (**Figure S1G**). Altogether, we found that all four RNA modifications studied (m^6^A, m^5^C, hm^5^C and Y) cause base-calling ‘errors’, and that these ‘errors’ follow specific patterns that depend on the RNA modification type.

### Y modifications can be detected *in vivo,* in the form of U-to-C mismatches and with single nucleotide resolution

We then examined whether the results obtained using *in vitro* transcribed constructs would be applicable to *in vivo* RNA sequences. To this end, total RNA from *S. cerevisiae* was poly(A)-tailed to allow for ligation between the RNA molecules and the commercial ONT adapters, and then prepared for direct RNA sequencing (see *Methods*). Visual inspection of the mapped reads revealed that our approach captured a high proportion of full-length rRNA molecules, with a high proportion of base-calling errors present in 25s and 18s rRNAs, as could be expected from sequences that are highly enriched in RNA modifications (**Figure 2A**). By contrast, 5s and 5.8s rRNAs did not show such base-calling errors, in agreement with their low level of modification.

**Figure 2.**
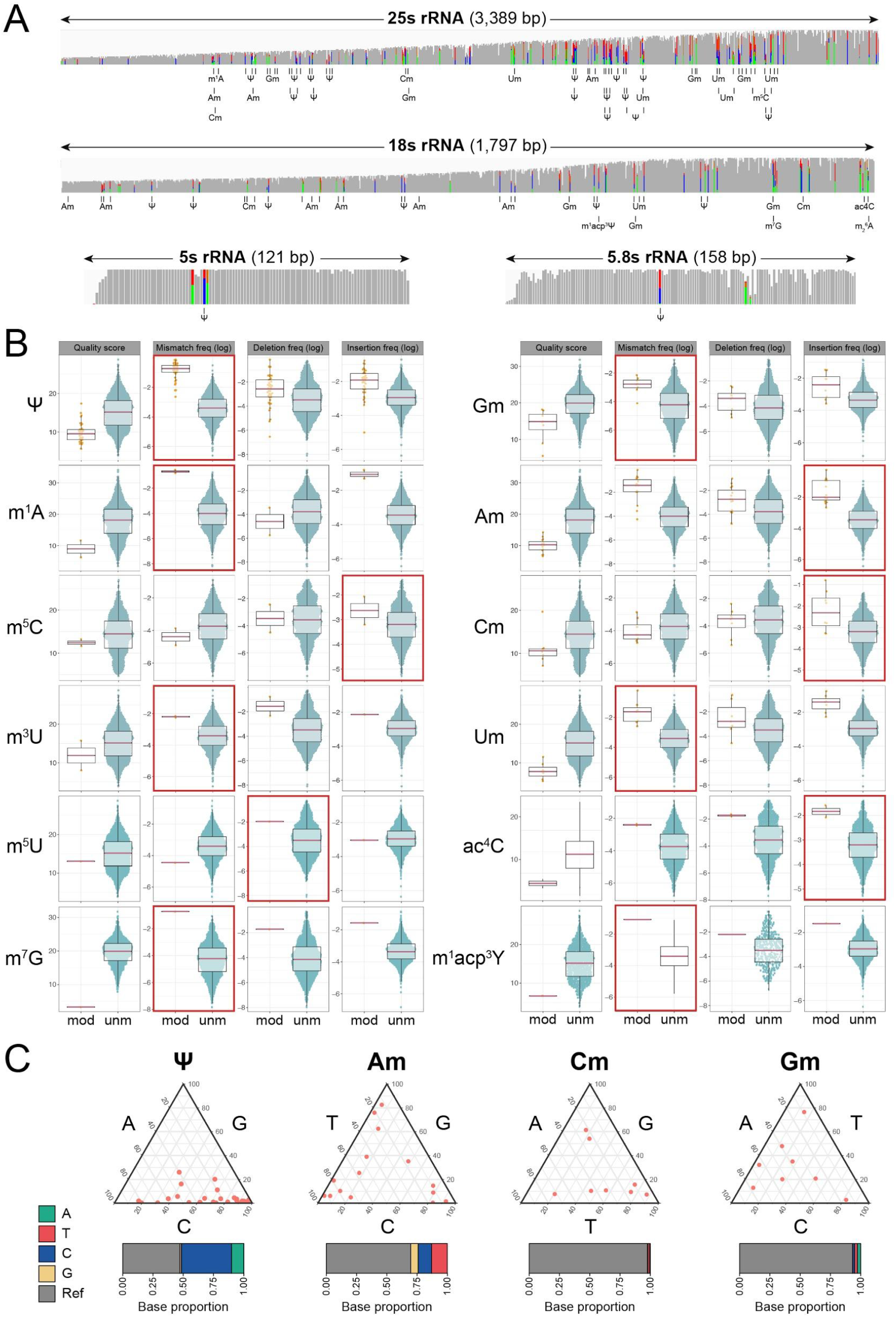
RNA modifications can be detected in yeast ribosomal RNA in the form of base-calling errors, and each RNA modification type shows a distinct ‘error’ signature. **(A)** IGV snapshots of yeast ribosomal subunits 5s, 5.8s, 18s and 25s. Known modification sites are indicated below each snapshot and nucleotides with mismatch frequencies greater than >0.1 have been colored and grey represents match to reference or no mismatch **(B)** Comparison of base-calling features (base quality, mismatch, deletion and insertion frequency) from distinct RNA modification types present in yeast ribosomal RNA. The most descriptive base-calling error per modification is outlined in red. Only RNA modification sites without additional neighboring RNA modifications in the 5-mer were included in the analysis: Y (n=37), Am (n=14), Cm (n=8), Gm (n=8), Um (n=7), ac^4^C (n=2), m^1^A (n=2), m^3^U (n=2), m^5^C (n=2), m^1^Acp^3^Y (n=1), m^5^U (n=1), m^7^G (n=1). Box, first to last quartiles; whiskers, 1.5x interquartile range; center line, median; dots: individual data points. (**C**) Ternary plots and barplots depicting the mismatch directionality for selected rRNA modifications (Y, Am, Cm, Gm). Y rRNA modifications tend towards U-to-C mismatches while Am, Cm and Gm modifications did not show specific mismatch directionality patterns. See also Figure S2 and S3.

Then, we systematically analyzed base-called features (mismatch, deletion, insertion and per-base qualities) in rRNAs, comparing the features from rRNA modified sites relative to unmodified ones (**Figure 2B**). We found that all rRNA modification types consistently led to decreased per-base qualities at modified sites, suggesting that per-base qualities can be employed to identify RNA modifications, but not the underlying RNA modification type. Moreover, we found that Y modifications caused significant variations in mismatch frequencies, in agreement with our observations using *in vitro* constructs. By contrast, other RNA modifications, such as 2’-O-methylcytidine (Cm) or 5-methylcytosine (m^5^C) did not appear in the form of increased mismatch frequencies at modified sites, but rather, in the form of increased insertions. In addition, Y modifications typically appeared in the form of U-to-C mismatches (**Figure 2C,** see also **Figure S2**), in agreement with our *in vitro* observations, whereas other RNA modifications such as 2’-O-methyladenosine (Am) did not cause mismatches with unique directionality. Thus, we conclude that distinct rRNA modification types can be detected in the form altered base-called features *in vivo*, and that their base-calling ‘error’ signature is dependent on the RNA modification type.

To confirm that the detected signal (U-to-C mismatches) in Y positions was caused by the presence of the Y modification, we compared ribosomal RNA modification profiles from wild type *S. cerevisiae* to those from snoRNA-knockout strains (snR3, snR34 and snR36), which lack Y modifications at known rRNA positions (**Figure 3A,** see also **Table S3**). Our results show that changes in rRNA modification profiles were consistently and exclusively observed in those positions reported as targets of each snoRNA. Moreover, the remaining Y-modified positions were not significantly altered by the lack of Y modifications guided by snR3, snR34 or snR36 (**Figure 3B**), suggesting that the modification status of Y sites is largely independent from other Y sites.

**Figure 3.**
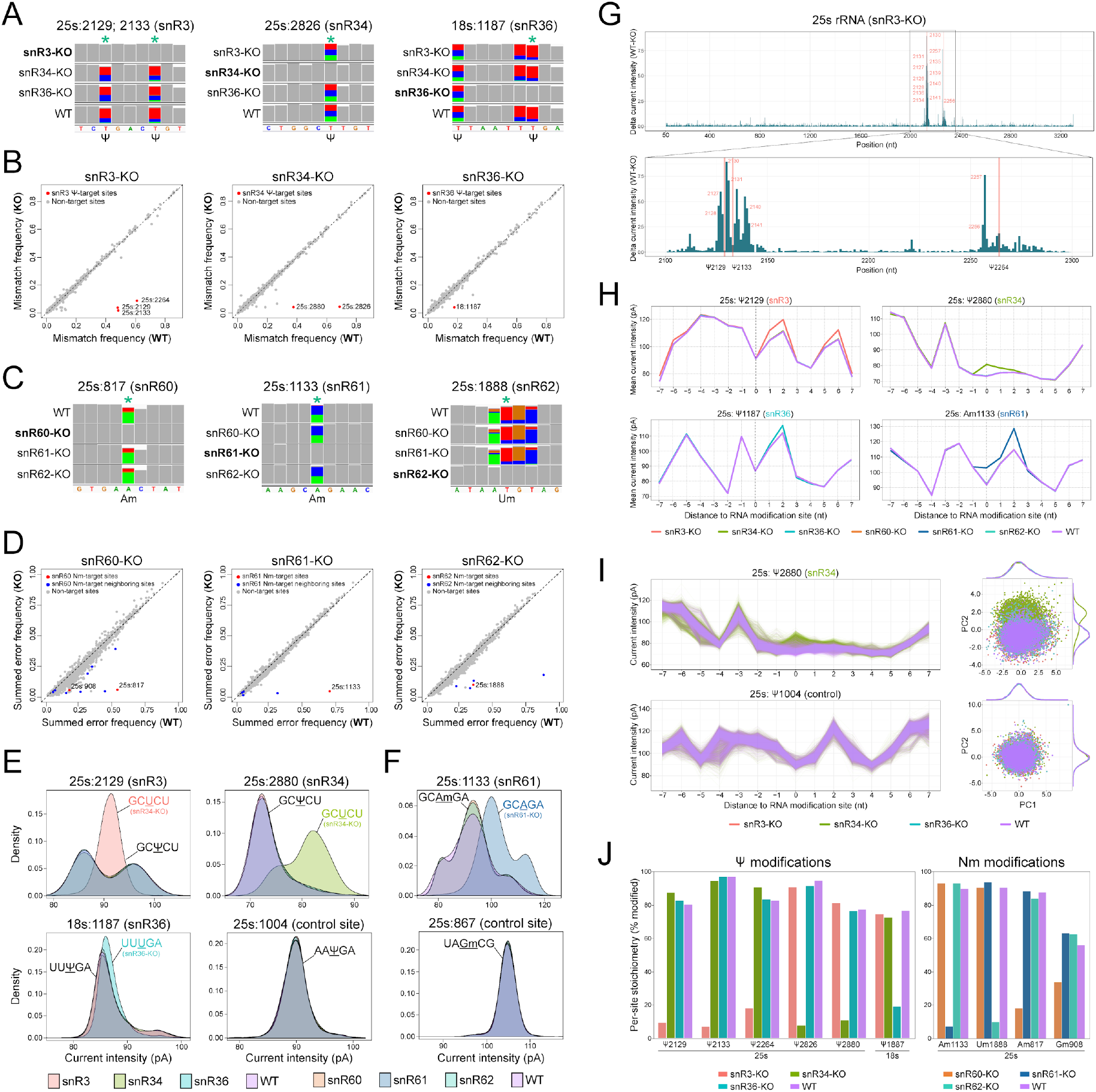
Pseudouridylation and 2’-O-methylations cause systematic base-calling ‘errors’ as well as altered current intensities, and their signature disappears upon depletion of snoRNAs guiding the modification. **(A)** IGV snapshots of wild type and three snoRNA-depleted strains depicting the site-specific loss of base-called errors at known Y target positions (indicated by asterisks). Nucleotides with mismatch frequencies greater than 0.1 have been colored. **(B)** Comparison of snoRNA knockout mismatch frequencies for each base, relative to wild type, with snoRNA targets sites indicated in red, and non-target sites in gray. **(C)** IGV snapshots of wild type and three snoRNA knockout yeast strains depicting the site-specific loss of base-calling errors at known Nm target positions. Nucleotides with mismatch frequencies greater than 0.1 have been colored. **(D)** Comparison of snoRNA knockout summed error frequencies for each base, relative to wild type, with snoRNA targets sites indicated in red, neighboring sites in blue and non-target sites in gray. **(E,F)** Distributions of per-read current intensity at known Y-modified (E), 2’-O-methylated (F) and negative control sites. Current intensities at Y and 2’-O-methylated positions were altered upon deletion of specific snoRNAs relative to wild type, whereas no shift was observed in control sites. **(G)** Current intensity changes along the 25s rRNA molecule upon snR3 depletion, relative to the wild type strain. In the lower panel, a zoomed subset focusing on the two regions with the most significant current intensity deviations is shown; the first one comprising the 25s:Y2129 and 25s:Y2133 sites, and the second one comprising the 25s:Y2264 site. **(H)** Comparison of current intensities in the 15-mer regions surrounding Y and 2’-O-methyl knockout sites, for each of the 4 strains. The dotted vertical line indicates the modified position. See also Figure S4 for current intensity changes in other knockout strains and sites. **(I)** Per-read current intensity analysis centered at the 25s:Y2880 site targeted by snR34 (upper panel) and a control site, 25s:Y2880, which is not targeted by any of the knockouts (lower panel). For each site, Principal Component Analysis was performed using 15-mer current intensity values, and the corresponding scatterplot of the two first principal components (PC1 and PC2) is shown on the right, using as input the same read populations as in the left panels. Each dot corresponds to a different read, and is colored according to the strain. **(J)** Predicted stoichiometry of Y- and Nm-modified sites using a k-nearest neighbors (KNN) algorithm trained to classify the reads into 2 classes: modified or unmodified. The features used to predict modifications status of every read from which stoichiometry was calculated were signal intensity (positions −1,0,+1) and trace (positions −1,0,+1). See also Figures S4 and S5.

### 2’-O-methylations can be detected *in vivo* in the form of systematic base-calling ‘errors’, but their signatures vary across sites

We then sequenced 3 additional *S. cerevisiae* strains depleted of snoRNAs (snR60, snR61 and snR62 knockouts) guiding 2’-O-methylation (Nm) at specific positions (**Table S3**). In contrast to Y modifications, we found that 2’-O-methylations often caused increased mismatch and deletion signatures not only at the modified position, but also at neighboring positions (**Figure 3C,** see also **Figure S3A**). These errors disappeared in the knockout strain, suggesting that neighboring base-calling errors were indeed caused by the 2’-O-methylation (**Figure 3C**). In contrast to Y modifications, which mainly affected mismatch frequency, we observed that Nm modifications often affected several base-called ‘error’ features (mismatch, insertion and deletion frequency) (**Figure S3B**). Thus, we reasoned that combining all three features might improve the signal-to-noise ratio for the detection of 2’-O-methylated sites (**Figure 3D**), and found that the combination of features led to improved detection of Nm-modified sites, relative to each individual feature. We should note that position 25s:Gm908 was poorly detected in both wild type and snoRNA-depleted strains (**Figure S3A,B**) regardless of the feature combination used, likely due to the sequence context in which the site is embedded -a homopolymeric GGGG sequence-, which is often troublesome for nanopore base-calling algorithms.

### Current intensity variations can be used to detect Y and Nm RNA modifications, but do not allow accurate prediction of the modified site

We then wondered whether Y and Nm sites would also be detected at the level of current intensity changes. We observed that certain Y and Nm-modified sites, such as 25s:Y2129 or 25s:Am1133, showed drastic alterations of their current intensity values in the snoRNA-depleted strain, while no significant alteration was observed in control sites (**Figure 3E,F**). However, the distribution of current intensities in some sites did not significantly change in the knockout strain (18s:Y1187, **Figure 3E** lower panel) or did not differ in their mean (25s:Y2133, **Figure S4A**).

We hypothesized that deviations in current intensity alterations might not always be maximal in the modified site, but might sometimes appear in neighbouring sites. To test this, we examined the difference in current intensity values along the rRNA molecules for each wild type-knockout pair (**Figure 3G**, see also **Figure S4B**). As expected, we found that the depletion of snR3 led to two regions with altered current intensity values along the 25s rRNA - one comprising the 25s:Y2129 and 25s:Y2133 sites, and the second comprising the 25s:Y2264 site. However, the highest deviations in current intensity were not observed at the modified site (**Figure 3G** lower panel). From all 6 Y sites that were depleted in the 3 knockout strains studied, only 2 of them (25s:Y2826 and 25s:Y2880) showed a maximal deviation in current intensity in the modified site (**Figure 3H,** see also **Figure S4C**). Similarly, depletion of Nm sites led to changes in current intensity values, but the largest deviations were not observed at the modified site. Thus, we conclude that current intensity-based methods can detect both Y and Nm RNA modifications; however, base-calling errors are a better choice to achieve single nucleotide resolution, at least in the case of Y RNA modifications.

### Per-read current intensity analysis of Y- and Nm-modified sites allows binning of individual reads based on their modification status

Direct RNA sequencing produces current intensity measurements for each individual native RNA molecule. Thus, native RNA sequencing can in principle estimate modification stoichiometries by identifying the proportion of reads with altered current intensity at a given site. To reveal whether current intensity alone would be sufficient to bin the reads into modified and unmodified populations, we first examined the per-read current intensity values of wild type and knockout strains at the Y- and Nm-depleted sites. We found that there was a significant variability across reads, even when 100% of the positions are unmodified, however, we were able to observe robust differences in current intensities across strains at the per-read level (**Figure 3I,** upper panel). As a control, we performed the same analysis in Y sites unaffected by snoRNA depletion, finding no differences between wild type and knockout strains at these positions (**Figure 3I,** lower panel). However, in some sites such as 18s:1187, the per-read shifts in current intensity between the wild type and knockout strain were far more modest (**Figure S4D**).

We then performed Principal Component Analysis (PCA) of the current intensity values corresponding to the 15-mer regions that contained the modified site, for all snoRNA-depleted strains affecting Y (snR3, snR34, snR36) and Nm modifications (snR60, snR61, snR62), as well as for the wild type strain (**Figure 3I** right panels, see also **Figure S4E**). As could be expected based on the per-read current intensity plots, we observed that the reads clustered into two distinct populations: the first cluster mainly comprised unmodified reads from the snoRNA-depleted strain, whereas the second comprised reads from the 3 other strains, which are mostly modified.

To our surprise, we observed that *Nanopolish* software did not resquiggle the reads evenly across sites. For example, it failed at resquiggling the majority of reads in the region surrounding 25s:Y2264 **(Figure S4D**). Thus, we examined whether the *Tombo* resquiggling algorithm, which uses global resquiggling instead of local resquiggling, might overcome this limitation, finding that *Tombo* resquiggling led to a global increase in the proportion of resquiggled reads (**Figure S5A**). Moreover, *Tombo* was equally effective at resquiggling both modified and unmodified reads, whereas *Nanopolish* preferentially resquiggled unmodified reads relative to modified ones, biasing the unmodified:modified proportion up to 7:1 (**Figure S5B**). This uneven resquiggling from *Nanopolish* implies that using *Nanopolish* for predicting RNA modification levels at individual sites may cause a dramatic bias in the predicted stoichiometry of individual sites, especially in scenarios where RNA modifications are substoichiometric, such as mRNAs. Thus, based on these results, we decided to adopt *Tombo* resquiggling instead of *Nanopolish* resquiggling for the prediction of RNA modification stoichiometries from individual RNA reads in all our downstream analyses.

### Stoichiometry prediction of Y and Nm-modified sites using signal intensity, dwell time and trace

Our results show that the presence of Y and Nm modifications lead to significant alterations in the current intensity profiles at the modified region (e.g. 25s:Y2880, **Figure 3H-I**). However, in other sites such as 18s:Y1187, current intensity alone was insufficient to bin the reads into two separate clusters (**Figure S4D,E**), suggesting that, in addition to current intensity, other features might be needed to distinguish modified from unmodified reads.

Previous works predicting DNA modifications from individual nanopore reads have typically relied on features such as signal intensity or dwell time to distinguish modified and unmodified read populations 46–49. Here, in addition to these two features, we explored whether the use of ‘trace’ (also termed ‘base probability’), which is reported directly by *Guppy* into the base-called FAST5 files, would improve our ability to predict RNA modification stoichiometry. To this end, we first examined how the presence of Y and Nm modifications altered each of the features (signal intensity, dwell time and trace) in Y and Nm modified sites by comparing the observed features in wild type and snoRNA-deficient strains, both at snoRNA-targeted positions and control sites (**Figure S6**). Our results show that in addition to signal intensity, base probability (trace) was significantly different in the snoRNA-deficient strains in all examined sites. Moreover, in some sites such as 25s:Y2264, trace was the most altered feature from those examined. By contrast, we found that dwell time was not consistently different in snoRNA-targeted sites relative to wild type (e.g. 25s:Y2264, 25s:Y2826, 18s:Y1187).

We then proceeded to systematically benchmark the use of distinct features for RNA modification stoichiometry. To this end, we built *nanoRMS*, a software that extracts the distinct features (signal intensity, trace and dwell time) from individual reads, and then predicts RNA modification stoichiometry by using distinct feature combinations as well as various machine learning algorithms. Firstly, we generated different mixes of modified (wild type) and unmodified (knockout) reads to simulate varying read stoichiometry (0, 20, 40, 60, 80 and 100%), for each of the Y and Nm positions for which knockouts were available (**Table S3**). Then, we examined how different supervised and unsupervised algorithms would predict the stoichiometry of each of the sites, and using distinct combinations of the 3 features (signal intensity, trace and dwell time) for each individual site (**Figure S5C**). Our results show that the combination of signal intensity and trace outperformed all the other feature combinations for predicting both Y and Nm modification stoichiometry, and that the supervised k-nearest neighbor (KNN) was the best performing algorithm. The k-means clustering algorithm (KMEANS) was the best-performing algorithm among the unsupervised clustering methods tested, although its the performance in predicting Y modification stoichiometry was slightly better than in the case of Nm modification stoichiometry predictions. Overall, we find that *nanoRMS* can accurately predict Y and Nm RNA modification stoichiometry from individual RNA reads (**Figure 3J**), with predicted stoichiometry values that are similar to those that have been previously reported by Mass Spectrometry ^50^ (**Table S4**).

### *De novo* prediction of Y modifications reveals a novel Pus4-dependent mitochondrial rRNA modification

The identification of RNA modification-specific signatures allows us to perform *de novo* prediction of Y RNA modifications transcriptome-wide using direct RNA sequencing. In this regard, *S. cerevisiae* mitochondrial rRNAs remains much less characterized than cytosolic rRNAs, with only 3 modified sites identified so far in *S.cerevisiae* LSU (21s) ^51^, and none in SSU (15s) rRNAs. Thus, we hypothesized that direct RNA might reveal previously uncharacterized Y-modified sites in mitochondrial rRNAs. To this end, we first determined the ‘error’-based thresholds (mismatch frequency and C mismatch frequency) that would distinguish unmodified uridines from pseudouridines in cytosolic rRNAs (**Figure 4A**). We then applied this filter to predict Y modifications on 15s rRNA and 21s rRNA, identifying two novel candidate Y sites (15s:854 and 15s:579) that displayed high modification frequency as well as U-to-C mismatch signature (**Figure 4B,C**).

**Figure 4.**
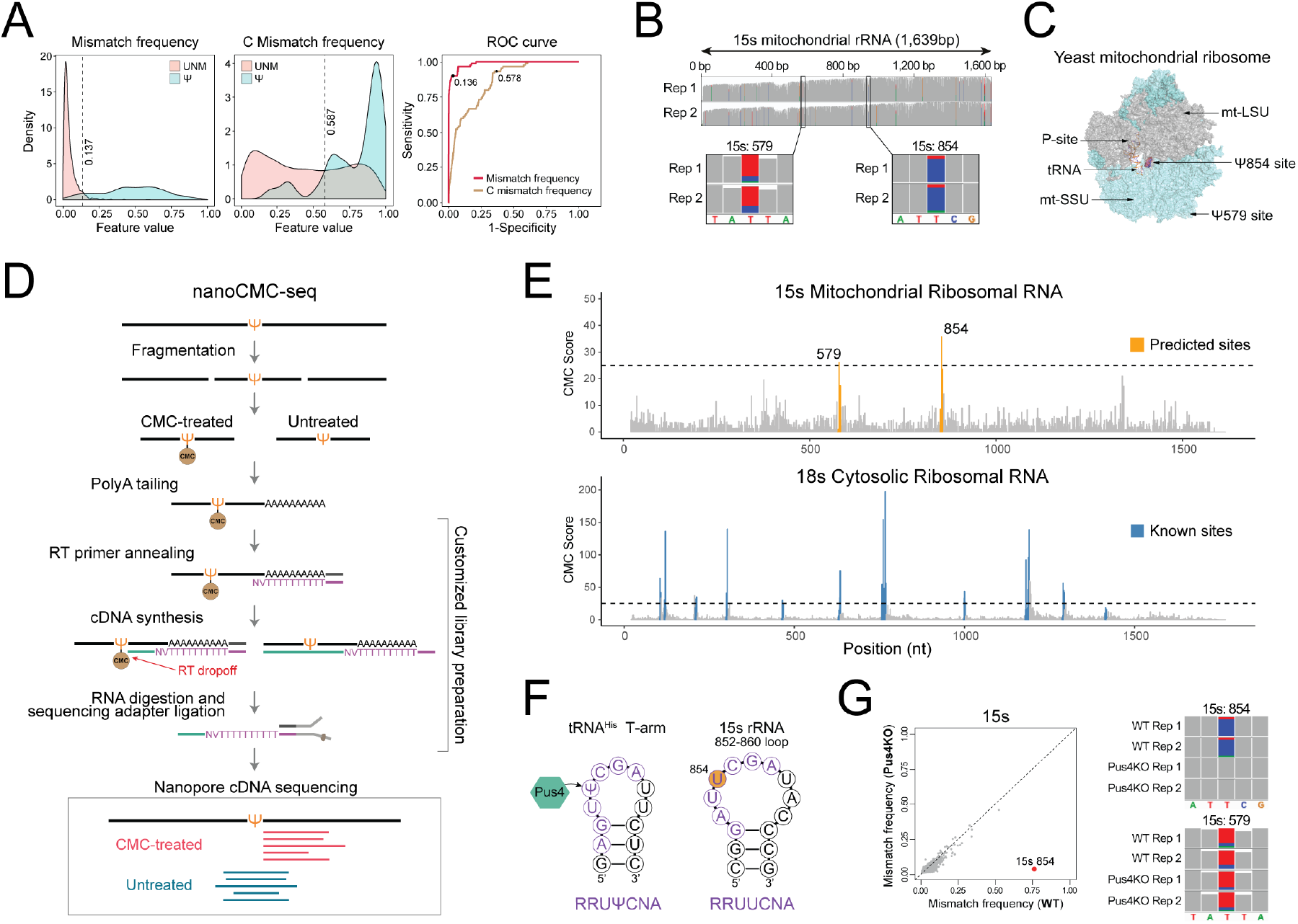
*De novo* prediction of Y modifications reveals a novel Pus4-dependent mitochondrial rRNA modification. **(A)** Density distributions of mismatch frequency and C mismatch frequency in unmodified uridine positions (red) and pseudouridine positions (cyan). The dashed lines represent the optimal cutpoints between two groups determined by maximizing the Youden-Index. In the right panel, the ROC curve illustrates the sensitivity and specificity at these two cutpoints. **(B)** IGV coverage tracks of the 15s mitochondrial rRNA, including a zoomed version showing the tracks centered at the 15s:854 and 15s:579 sites, in two biological replicates. Nucleotides with mismatch frequencies greater than 0.15 have been colored. **(C)** Location of the putative Y854 modified site in the yeast mitochondrial ribosome. The LSU has been colored in cyan, whereas the SSU has been colored in gray. The tRNA is located in the P-site of the ribosome. The PDB structure shown corresponds to 5MRC. **(D)** Validation of the 15s:579 and 15s:Y854 with nanoCMC-Seq, which combines CMC treatment with Nanopore cDNA sequencing in order to capture RT-drops that occur at Y-modified sites upon CMC probing. RT-drops are defined by counting the number of reads ending (3’) at a given position. CMC-probed samples will cause accumulation of reads with the same 3’ ends at positions neighboring the Y site (red), whereas untreated samples will show random distribution of 3’ ends of their reads (teal). **(E)** Predicted Y sites U854 and U579 (orange) in the 15s rRNA are validated using nanoCMC-seq (upper panel). Dashed lines indicate the CMC-score threshold used for determining the positive sites (upper panel). As a control, we analyzed the nanoCMC-seq results in other rRNAs (lower panel), finding that all positions with a significant CMC Score (>25) correspond to known Y rRNA modification sites (blue). See also Figure S7A for CMC scores in additional rRNA transcripts. **(F)** The candidate Y854 site is located at the 852-860 loop of the 15s rRNA, which resembles the t-arm of the tRNAs that is modified by Pus4. The binding motif of Pus4 (RRUUCNA) matches the motif surrounding the 854U site ^4^. **(G)** Scatterplot of mismatch frequencies in WT and Pus4KO cells, showing that the only significant position affected by the knockout of Pus4 is 15s:U854 (left panel). IGV coverage tracks showing that Pus4 knockout leads to depletion of the mismatch signature in the 15s:854 position (right panel), but not at the 15s:579 position.

To further confirm that the two predicted 15s rRNA sites are pseudouridylated, we developed nanoCMC-seq, a novel protocol that identifies Y modifications by coupling CMC probing with nanopore cDNA sequencing. This method allows capturing reverse-transcription drop-off information by sequencing only the first-strand cDNA molecules of CMC-probed RNAs using a customized direct cDNA sequencing protocol (**Figure 4D,** see also *Methods*). We found that NanoCMC-seq captured known sites in cytoplasmic rRNA with a very high signal-to-noise ratio, as well as confirmed the existence of Y in position 854 and 579 of 15s rRNA, validating our *de novo* predictions using direct RNA sequencing (**Figure 4E,** see also **Figure S7A**).

We then examined the sequence context of these two novel 15s rRNA modifications. We observed that 15s:Y854 was embedded in a similar sequence context and structure as the t-arm of tRNAs, which contains a pseudouridylated (Y55) position placed by Pus4 (**Figure 4F**). Given the resemblance between these two sequences and structures, we hypothesized that Pus4 might be responsible for this modification. To validate our hypothesis, we sequenced total RNA from a *S. cerevisiae* Pus4 knockout strain, finding that the 15s:854 position loses its mismatch signature upon deleting Pus4 gene without altering the base-called feature of any other position on the ribosomal RNAs, confirming that not only this site is pseudouridylated, but also that it is Pus4-dependent (**Figure 4G,** see also **Figure S7B)**. Additionally, we observed that previously reported Pus4 target sites (TEF1:239,TEF2:239) ^3–5^ completely lost their mismatch signature in Pus4 knockout cells **(Figure S7B,C)**, confirming that our method is able to capture previously reported Pus4-dependent Y sites, in addition to novel ones.

### rRNA modification profiles do not vary upon exposure to oxidative or thermal stress, whereas Y modification levels in several snRNAs and snoRNAs significantly change upon heat stress

Ribosomal RNAs are extensively modified as part of their normal maturation, and their modification landscape is relatively well-defined for a series of organisms ^38,52–55^. Typically placed by either stand-alone enzymes or snoRNA-guided mechanisms, rRNA modifications tend to cluster in functionally important sites of the ribosome, stabilizing its structure and fine-tuning its decoding capacities ^56^. Despite the central role that rRNA molecules play in protein translation, recent evidence has shown that rRNA modifications are in fact dynamically regulated ^57,58^, and that their alterations can lead to disease states ^40,41,59–65^. Moreover, it has been shown that some pseudouridylated and 2′-O-methylated rRNA sites are only partially modified, and that their stoichiometry is cell-type dependent, suggesting that rRNAs modifications may be an important source of ribosomal heterogeneity ^42,50,53,66–68^. However, a systematic and comprehensive analysis of which environmental cues may lead to changes in rRNA modification stoichiometries, which RNA modifications may be subject to this tuning, and to which extent, is largely missing.

To assess whether rRNA modification profiles change in response to environmental stimuli, we treated *S. cerevisiae* cells with diverse environmental cues (oxidative, cold and heat stress) and sequenced their RNA, in biological duplicates, using direct RNA sequencing. Firstly, we examined the reproducibility across biological replicates, finding that the rRNA modification profiles from independent biological replicates were highly reproducible (pearson r2=0.976-0.996). Then, we examined whether exposure to stress (oxidative, cold and heat stress) would lead to significant changes in base-calling ‘errors’ in rRNA molecules, finding no significant differences in rRNA modification profiles between normal and stress conditions (**Figure 5A**). By contrast, we recapitulated previously reported changes in snRNA Y modifications upon exposure to environmental cues^4^ (**Figure 5B,** see also **Figure S7D**), as well as identified 8 additional Y modification sites in snRNAs and snoRNAs whose stoichiometry varies upon heat exposure, which had not been previously described (**Figure 5,** see also **Figure S7E and Table S5**) ^3,4,37,69^. Overall, our approach confirmed previous reports and predicted novel Y sites in ncRNAs whose modification levels vary upon heat shock exposure (**Figure 5B-D,** see also **S7D-E**), but did not identify any rRNA modified site to be varying in its stoichiometry upon any of the tested stress conditions.

**Figure 5.**
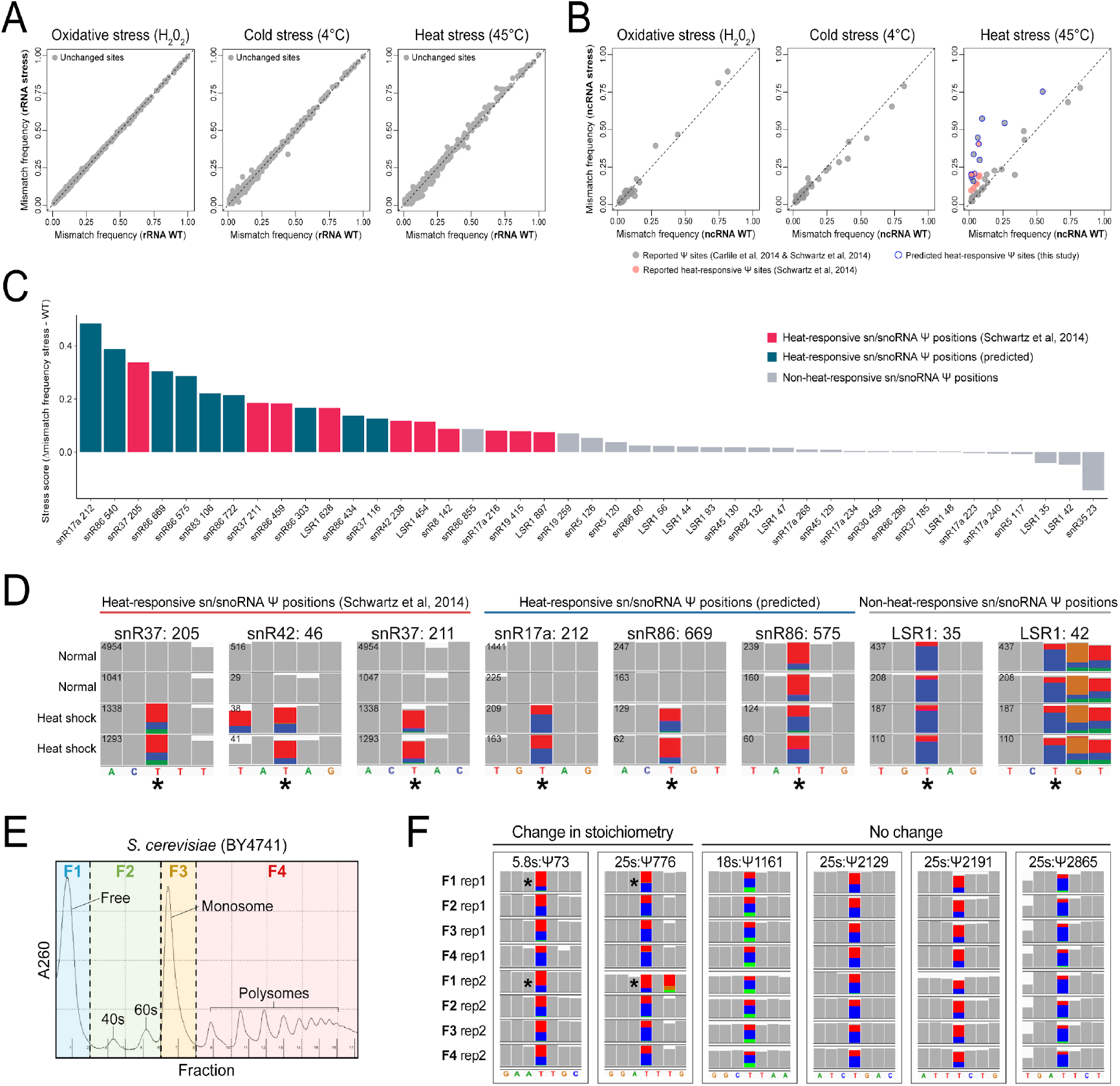
Comparative analysis of yeast rRNA and snRNA Y modifications upon distinct environmental stresses identifies previously known and novel heat-sensitive snRNA and snoRNA Y modifications. **(A)** Comparison of mismatch frequencies for all rRNA bases from untreated or yeast exposed to oxidative stress (H_2_O_2_, left panel), cold stress (4°C, middle panel) or heat stress (45°C, right panel). Each dot represents a uridine base. All rRNA bases from cytosolic rRNAs were included in the analyses. **(B)** Comparison of mismatch frequencies in untreated versus stressed-exposed yeast cells (oxidative, cold or heat), in previously reported ncRNA Y sites ^3,4^. **(C)** Stress scores in sn/snoRNA Y sites calculated by Δ mismatch frequency between heat shock and WT. **(D)** IGV snapshots of normal condition (rep1 and rep2) and heat shock condition (rep1 and rep2) yeast cells zoomed into the known sn/snoRNA Y positions (indicated by an asterisk). Nucleotides with mismatch frequencies greater than 0.1 have been colored. Coverage for each position/condition is given on the top left of each row. **(E)** Profiles of ribosomal fractions isolated from yeast grown under normal conditions, using sucrose gradient fractionation, including free rRNAs which are not assembled into ribosomal subunits (F1), rRNAs from 40s and 60s subunits (F2), rRNAs extracted from monosomal fractions (F3) and polysome fractions (F4). **(F)** IGV snapshots of the two Y sites that change stoichiometry between translational fractions and four representative Y sites that show no significant change. Nucleotides with mismatch frequencies greater than 0.1 have been colored. See also Figure S7.

### rRNA modification profiles do not vary across translational repertoires

Next we questioned whether pseudouridylation changes in distinct translational repertoires may be more nuanced, in that Y levels may differ between rRNAs present in different translational fractions along a polysome gradient, which would not be detected when examining rRNAs as a whole. To test this, we sequenced both total (input) and polysomal rRNAs from untreated and H_2_O_2_-treated yeast cells (**Figure S7F**). However, we observed no significant changes in Y rRNA modification profiles when comparing rRNAs from actively translating ribosomes in untreated versus H_2_O_2_-treated cells (**Figure S7G**).

In an attempt to further dissect the different translational repertoires into a higher number of rRNA pools, we sequenced: i) rRNAs from unassembled free rRNA fractions (F1), ii) rRNAs from 40s and 60s subunits (F2), iii) rRNAs from monosomal fractions (F3) and iv) rRNAs from polysomal fractions (F4) (**Figure 5E**). While two positions showed slightly decreased levels of Y (5.8s:Y73 and 25s:Y776) in the free rRNA fraction (F1) compared to assembled ribosomes and/or subunits, no significant changes were observed across the other translational fractions (**Figure 5F,** see also **Figure S7H**). Globally, these results indicate that differential rRNA modification is likely not a mechanism employed by yeast cells to adapt to environmental stress conditions, in agreement with previous observations ^3^.

### *De novo* prediction of Y modifications in mRNAs using direct RNA sequencing reveals novel Y sites that are Pus1, Pus4 and heat stress-dependent

Ribosomal RNAs are modified at very high stoichiometries ^50,53^. By contrast, other RNA molecules such as mRNAs are considered to be modified at much lower stoichiometries, making the detection of their RNA modifications a much more challenging task ^21^. To ascertain whether our methodology would be applicable to lowly modified RNA sites, such as those present in mRNAs, we first assessed the performance of *nanoRMS* in RNA molecules that contained Y RNA modifications at low RNA modification stoichiometries (0, 3, 7 and 20%) (**Figure 6A**, see also *Methods*). These synthetic RNA molecules were produced by *in vitro* transcription, and their relative incorporation of Y RNA modifications was validated using Mass Spectrometry. We then examined the quantitative performance of *nanoRMS* under low stoichiometry conditions using both KNN and k-means, finding that the combination of signal intensity and trace features yielded the most accurate results in terms of stoichiometry prediction (**Figure 6B**), in agreement with our previous results (**Figure S5C**).

**Figure 6.**
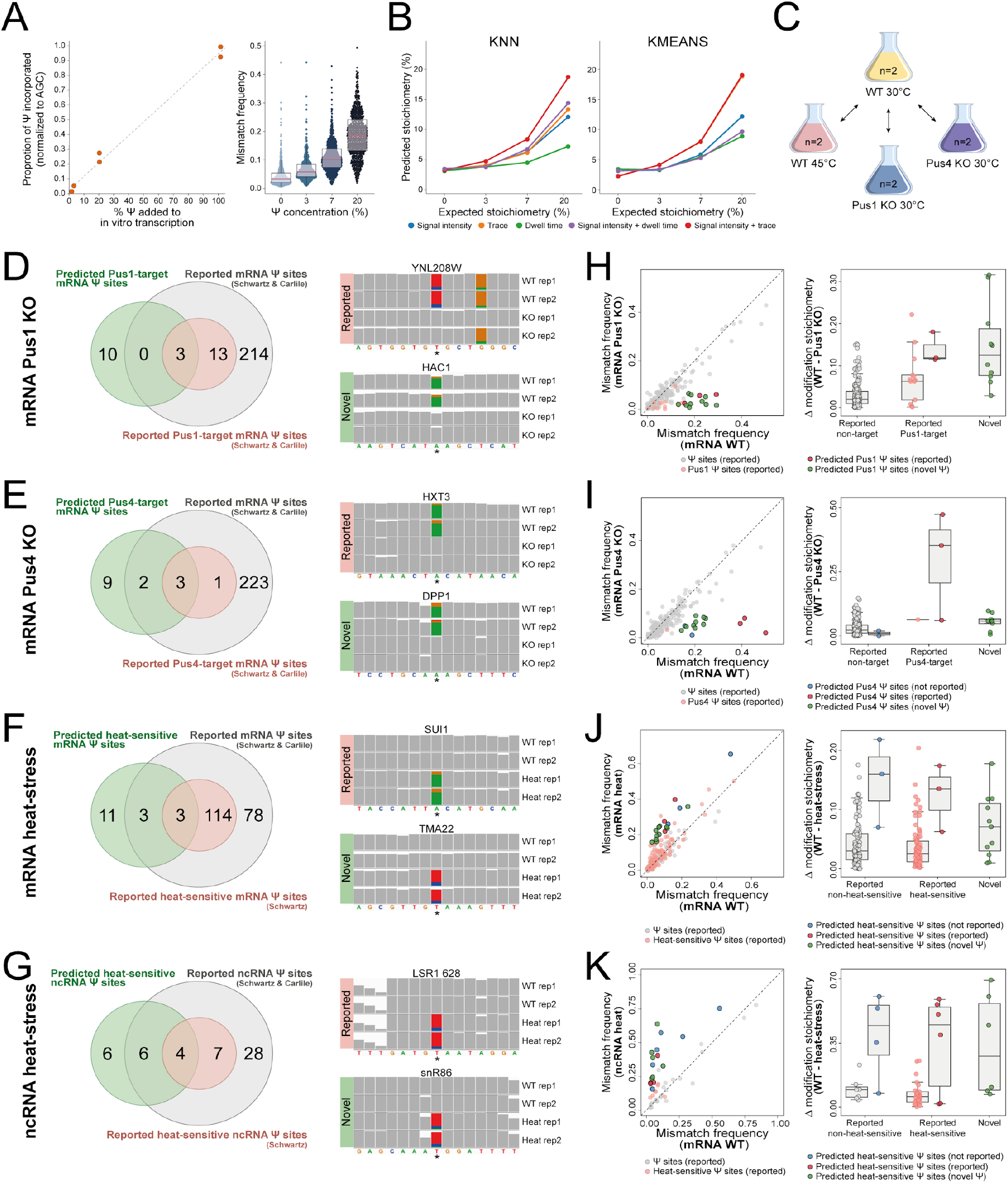
Quantitative prediction of pseudouridine stoichiometry transcriptome-wide and systematic benchmarking of *nanoRMS* using RNA molecules with diverse modification stoichiometries. **(A)** LC-MS validation of pseudouridine incorporation at different proportions (0%, 3%, 20%, 100%) in the *in vitro* transcribed products, relative to the expected incorporation (% YTP relative to UTP) (left panel). In the right panel, the dotplot illustrates the mismatch frequency distribution of the uridine positions in the *in vitro* transcribed products incorporated with different concentrations of Y. Each dot represents one uridine position. **(B)** Stoichiometry predictions of the Y incorporated *in vitro* transcription products using two different algorithms (KNN and k-means) with different current information (middle right and right panels). **(C**) Conditions and strains used to predict Y mRNA modifications transcriptome-wide. (**D-K)** Transcriptome-wide Y RNA modification predictions and predicted stoichiometries in mRNAs and ncRNAs, for Pus1-dependent mRNA Y sites (D,H), Pus4-dependent mRNA Y sites (E,I), heat stress-dependent mRNA Y sites (F,J) and heat stress-dependent ncRNA Y sites (G,K). (**D-G**) Venn diagrams depict the overlap between Y sites predicted by our analysis and the previously reported pseudouridine sites. IGV snapshots of reported and novel predicted sites illustrate the absence of the mismatch signature in the Pus1 (D) or Pus4 (E) knockout samples as well as under normal conditions, relative to heat stress conditions in mRNA (F) and ncRNA (G) are also shown. The reported or predicted Y site is indicated by an asterisk. Nucleotides with mismatch frequencies greater than 0.15 have been colored. We should note that IGV snapshots that show a reference “A” with mismatch signature to G are genes that are in the minus strand (and thus are in reality positions showing U-to-C mismatch signatures). (**H-K**) Quantitative analysis of previously reported and *de novo* predicted Y sites in mRNAs and ncRNAs. In the left panels, comparative scatterplots of mismatch frequency illustrate differentially modified sites of reported and *de novo* predicted Y sites. In the right panels, stoichiometry prediction differences between WT and knockout strains (H-I) or between normal and heat stress conditions (J-K) are depicted in the form of boxplots. Each dot represents a Y site.

Next, we sequenced polyA(+)-selected RNA from *S. cerevisiae* wild type, Pus1 knockout, Pus4 knockout and heat stress-exposed strains using direct RNA sequencing, in biological duplicates. Considering that mRNA sites are lowly modified, we restricted our *de novo* identification of mRNA Y sites to those whose base-calling ‘error’ features significantly changed between pairwise conditions (**Figure 6C**, see also *Methods*), met the pseudouridine ‘error’ signature, and had a minimum coverage of 30 reads in both conditions and biological replicates (**Table S6,** see also *Methods*). Through this approach, we predicted 13 Pus1-dependent Y mRNA modifications, 14 Pus4-dependent Y mRNA modifications, 17 heat stress-dependent Y mRNA modifications and 16 heat stress-dependent Y ncRNA modifications, respectively (**Figure 6D-G** left panels, see also **Tables S7-10**), some of which were not previously reported to be Y-modified.

*NanoRMS* recovered 11% of previously reported Pus1-dependent Y sites as well as 75% Pus4-dependent Y sites, in addition to predicting 10 novel Pus1 and 11 novel Pus4-dependent mRNA Y-modified sites (**Table S7** and **S8**). These novel predicted Y mRNA sites displayed similar mismatch signatures to those observed in previously reported Y sites (**Figure 6D-E**, right panels), were highly replicable across biological replicates, and their signature disappeared in Pus1 or Pus4 knockout strains. Similarly, *nanoRMS* was able to capture previously reported heat-responsive Y sites present in mRNAs and ncRNAs, which resulted in predicting 17 heat-responsive Y mRNAs sites, among which 6 of them were previously reported Y sites (**Figure 6F**, see also Table S9), as well as 16 heat-responsive Y ncRNAs sites, from which 10 were previously reported Y sites (**Figure 6G**, see also **Table S10**).

Surprised by the relatively poor overlap between our predictions and previously reported Pus1 mRNA Y-modified sites (3 out of 16 sites), as well as poor overlap between predicted and previously reported heat stress-dependent sites (7 out of 128 sites), we inspected the individual per-read features at previously reported Pus1- and heat stress-dependent sites (**Figure S8A,B)**. Indeed, the Y sites that *nanoRMS* did not report as Pus1 or heat stress-dependent were not significantly different for any of the features examined (current intensity, dwell time or trace). Thus, we wondered whether some of these sites might have been misassigned as Pus1 or heat stress-dependent by previous works. Indeed, if we examine the overlap between mRNA and ncRNA Y sites predicted by the two previously published studies using CMC probing coupled to Illumina sequencing ^3,4^, which we used to define the set of ‘previously reported Pus1-, Pus4- and heat stress-dependent Y sites’, we observed that the overlap between the two studies was in fact very poor (**Figure S8C**), both when examining the set of predicted mRNA and ncRNA Y sites (7% and 17%, respectively), as well as when examining the sets of predicted Pus1- and Pus4-dependent mRNA and ncRNA Y sites (6% and 50%, respectively). Altogether, our approach detected 100% of Pus1- and Pus4-dependent sites that were identified by both studies, but very few of those that were identified by only one of the studies. Thus, we conclude that the poor overlap between our results and previously reported Y sites is in fact a direct consequence of the poor overlap between the set of predicted Pus1-, Pus4- and heat stress-dependent mRNA and ncRNA Y sites by the two previous studies (**Figure S8C**).

Finally, we applied *nanoRMS* to predict the modification stoichiometry of all previously reported and novel Y sites predicted in mRNAs and ncRNAs. To this end, reads were classified based on the per-read signal intensity and trace features from positions −1, 0, and +1 using the k-means unsupervised clustering algorithm (**Figure 6H-K**). As expected, we observed that per-read stoichiometry predictions were low in non-targeted Y sites. By contrast, predicted Y Pus1/Pus4/heat stress-dependent sites (which included both previously reported and novel Y sites) typically showed significant RNA modification stoichiometry changes, ranging from 5 to 50% change in their Y modification stoichiometries between the two conditions.

Altogether, our results show that the use or differential ‘error’ Y signatures are a useful approach to identify dynamic Y RNA modifications across two conditions even at low stoichiometry sites, and that *nanoRMS* can be used to *de novo* predict and quantify the RNA modification stoichiometry dynamics at these sites from their per-read features, both in previously reported Y sites, as well as in *de novo* predicted Y sites.

## DISCUSSION

RNA modifications are key regulators of a wide range of biological processes ^70–72^. They can modulate the fate of RNA molecules, such as mRNA splicing ^73–75^ or mRNA decay ^76,77^, as well as affect major cell and organism-level decisions, such as cellular differentiation ^78,79^ and sex determination ^18,80,81^. While the biological relevance of RNA modifications is out of question, a major difficulty in studying them has been the need for tailored protocols to map each modification individually ^20,82^. In this context, direct RNA nanopore sequencing has emerged as a promising platform that can overcome many of the limitations that NGS-based methods suffer from, as it can sequence full-length native RNA molecules, including their RNA modifications.

In the last few years, direct RNA nanopore sequencing has successfully been applied to reveal long-read native RNA transcriptomes from a wide variety of organisms ^29–31,83–86^. However, the detection of distinct RNA modification types in individual native RNA molecules is still an unsolved challenge. The ideal solution would be that direct RNA base-calling algorithms, such as *Guppy* or *Albacore*, would predict RNA modifications on-the-fly during the base-calling step, in a similar fashion to what *Guppy* 3.5.+ and later versions do in genomic DNA runs, where the base-calling algorithm can identify 6 different DNA nucleotides within the reads: A, G, C, T, m^6^A and m^5^C. However, this is not yet the reality for direct RNA sequencing, partly due to the higher noise-to-signal ratio of RNA nanopore reads. Consequently, solutions to identify RNA modifications in direct RNA sequencing data have so far relied on the use of post-processing software ^28–30,32,33,87^.

While both current intensity-based and ‘error’-based methods have proven useful strategies to detect RNA modifications, these methods have been mainly focused on the detection of m^6^A ^29–31,33^-, and are typically unable to predict which RNA modification type they are in fact detecting (e.g. m^6^A, Y, Am or m^5^C) ^28,49^. Moreover, current algorithms to study RNA modifications using direct RNA sequencing are not quantitative. To overcome these limitations, here we first explored how distinct RNA modifications may differentially affect direct RNA nanopore signals and base-calling ‘errors’. We find that different RNA modification types (e.g. Y versus m^5^C) produce distinct yet characteristic base-calling ‘error’ signatures, both *in vitro* (**Figure 1E-G, S1F**) as well as *in vivo* (**Figure 2**). Consequently, base-calling errors can be used not only to predict whether a given site is modified or not, but also to identify the underlying RNA modification type. While we should note that base-calling signatures depend to some extent on the surrounding sequence context, we find that Y modifications lead to robust U-to-C mismatch signatures, which can be exploited for *de novo* prediction of Y modifications (**Figure 4**). Through this approach, we identified two novel Y modifications in yeast 15s mitochondrial rRNA (15s:579 and 15s:Y854) that were not reported to date, as well as confirmed previously reported Y-modified sites in rRNAs, snRNAs and mRNAs (**Figures 3–6**). Moreover, we revealed that Pus4, which was previously thought to modify only tRNAs and mRNAs, is the enzyme responsible for placing Y854 in mitochondrial rRNA. These findings were further validated using nanoCMC-seq, a novel orthogonal method that can detect Y modifications with single nucleotide resolution by coupling CMC probing to nanopore cDNA sequencing (**Figure 4D**).

While we find that Y modifications can be detected both in the form of base-calling ‘errors’ and altered current intensities (**Figures 3**), we observe that the latter does not provide single nucleotide resolution, with maximal current intensity shifts are often seen a few nucleotides away from the real modified site, and that these shifts will also depend on the resquiggling algorithm used. Thus, current intensity-based methods alone may suffer from imprecisions in the assignment of the RNA-modified site. Here we propose that the combination of both approaches, i.e. base-called features and current intensity/trace features, is the optimal design to obtain stoichiometric information of Y-modified sites with single nucleotide resolution. Specifically, we show that once the site has been located using base-calling error features, per-read features (current intensity, trace and dwell time) from the regions surrounding Y or Nm-modified site are sufficient to robustly bin the reads into two separate clusters (modified and unmodified), and provide good estimates of Y and Nm modification stoichiometries (**Figure 3J** and **6B**).

One surprising feature of base-calling ‘errors’ is that fully modified sites do not always lead to same mismatch frequencies, suggesting that mismatch frequencies alone cannot be used per se as an estimation of the stoichiometry of the site (**Figure 2B**). While it is true that within the same sequence context, higher mismatch frequencies correspond to higher modification levels, this same rule cannot be used to compare across distinct RNA-modified sites. We speculate that the differences observed in mismatch frequency across different sites might be in fact a consequence of the deviation in current intensity of the modified k-mer relative to unmodified counterparts. For example, in the case of Y, the current intensity distribution of the Y-centered k-mers is shifted towards C-centered k-mers, and consequently, leading to U-to-C mismatch signatures (**Figure S8D**). However, the shift in current intensity may vary depending on the sequence context, leading to differences in mismatch frequencies across Y-modified sites (e.g. 25s:Y2826 compared to 25s:Y2880), despite having similar modification stoichiometries ^50^.

Finally, we should note that while *nanoRMS* allows predicting and studying the dynamics of diverse RNA modifications in a quantitative manner, there are caveats and limitations, leaving ample room for future improvements. First, not all RNA modifications lead to strong alterations in the base-calling features and/or current intensity patterns, such as 2’-O-methylcytosine (Cm), which is poorly detected in direct RNA sequencing datasets, compared to other RNA modifications (**Figure 2C**). Newer versions of protein nanopores, which are actively being developed, might lead to increased differences in current intensities when these RNA modifications pass through the nanopores. Second, the detection of RNA modifications is partly dependent on the sequence context; for example, we were unable to detect 25s:Gm908 (**Figure S3**). Similarly, some Y-modified sites, such as 18s:Y1187, cause weaker alterations in base-calling features and current intensity shifts than other Y-modified positions (**Figure 3**), although this limitation can be alleviated by the incorporation of additional features into the model (**Figure S5C**). Third, not all RNA modifications lead to base-calling errors with single nucleotide resolution, as with pseudouridine. For example, 2’-O-methylations often affect neighboring bases (**Figure 3C** and **S4A**), making it challenging to *de novo* predict modified sites without any prior information. Fourth, stoichiometry prediction is heavily affected by the choice of resquiggling algorithms (**Figure S5**). For example, we were unable to predict stoichiometry in 25s:Y2264 when using resquiggling due to the low number of reads that the *Nanopolish* algorithm was able to resquiggle (**Figure S4E**); however, this limitation could be overcome when using *Tombo* resquiggling, leading to stoichiometry predictions similar to those observed using Mass Spectrometry (**Figure 3J**). Future algorithms that improve the current intensity-to-base relationship will likely maximize our ability to extract modification information from direct RNA nanopore sequencing datasets. Finally, we should note that while *nanoRMS* was successful at detecting RNA modification stoichiometry changes as low as 5-10% (**Figure 6**), the detection of RNA modification changes in sites that show low modification stoichiometry was only possible when using comparison of pairwise conditions.

Despite these challenges and limitations, our work provides a novel framework for the systematic and comprehensive analysis of the epitranscriptome with single molecule resolution, showing that direct RNA sequencing can be employed to estimate Y and Nm modification stoichiometry as well as to *de novo* predict Y RNA modifications transcriptome-wide, in rRNAs, ncRNAs and mRNAs. Future work will be needed to functionally dissect the biological roles and dynamics of RNA modifications across further biological conditions and in disease states, to better comprehend how and when the epitranscriptome is tuned to regulate diverse cellular functions.

## ONLINE METHODS

### Yeast culturing

*Saccharomyces cerevisiae* (strain BY4741) was grown at 30°C in standard YPD medium (1% yeast extract, 2% Bacto Peptone and 2% dextrose). The deletion strains snR3Δ, snR34Δ and snR36Δ were generated on the background of the BY4741 strain by replacing the genomic snoRNA sequence with a *kanMX4* cassette as detailed in Parker et al. ^91^. Cells were then quickly transferred into 50 mL pre-chilled falcon tubes, and centrifuged for 5 minutes at 3,000 g in a 4°C pre-chilled centrifuge. Supernatant was discarded, and cells were flash frozen. For thermal stress, *Saccharomyces cerevisiae* BY4741 cultures were grown in 4 mL of YPD overnight at 30°C. The next day, cultures were diluted to 0.0001 OD600 in 200 mL of YPD and grown overnight at 30°C shaking (250 rpm). When the cultures reached an OD600 of 0.4-0.5, the cultures were divided into 3 × 50 mL subcultures, which were then incubated at 30°C (control), 45°C (heat shock) or 4°C (cold shock) for 1 hour. Cells were collected by pelleting and snap freezing. For the analysis of rRNAs modifications across polysomal fractions, yeast BY4741 starter cultures were grown in 6 mL YPD medium at 30°C with shaking (250 rpm) overnight. 100 mL of fresh YPD medium was inoculated with 10 μL of the stationary culture in a 250 mL erlenmeyer flask, in biological duplicates. Cells were incubated at 30°C with shaking (250 rpm) until the cultures reached mid-exponential growth phase (O.D_660_.~ 0.4-0.6). Yeast cells were then treated with 1 mM H_2_0_2_ or left without treatment (control) for 30 minutes. 1 mL of cycloheximide stock solution (10 mg/mL) was added to each culture. Pus4 knockout strains (BY4741 MATa pus4::KAN) and its parental strain were obtained from the Yeast Knockout Collection (Dharmacon) and grown under standard conditions in YPD (1% [w/v] yeast extract, 2% [w/v] peptone supplemented with 2% glucose) at 30°C unless stated otherwise.

### Total RNA extraction from yeast cultures

*Saccharomyces cerevisiae* BY4741 cells (strains: snR3Δ, snR34Δ snR36Δ, snR60Δ, snR61Δ, snR62Δ and WT) were harvested via centrifugation at 3000 rpm for 1 minute, followed by two washes with water. RNA was purified from pelleted cells using a MasterPure Yeast RNA extraction kit (Lucigen, MPY03100), according to manufacturer’s instructions. Total RNA was then treated with Turbo DNase (Thermo, #AM2238) with a subsequent RNAClean XP bead cleanup prior to starting the library preparation. For stress conditions and the Pus4KO strain, flash frozen pellets were resuspended in 700 μL Trizol with 350 μL acid washed and autoclaved glass beads (425-600 μm, Sigma G8772). The cells were disrupted using a vortex on top speed for 7 cycles of 15 seconds (the samples were chilled on ice for 30 seconds between cycles). Afterwards, the samples were incubated at room temperature for 5 minutes and 200 μL chloroform was added. After briefly vortexing the suspension, the samples were incubated for 5 minutes at room temperature. Then they were centrifuged at 14,000 g for 15 minutes at 4°C and the upper aqueous phase was transferred to a new tube. RNA was precipitated with 2X volume Molecular Grade Absolute ethanol and 0.1X volume Sodium Acetate. The samples were then incubated for 1 hour at −20°C and centrifuged at 14,000 g for 15 minutes at 4°C. The pellet was then washed with 70% ethanol and resuspended with nuclease-free water after air drying for 5 minutes on the benchtop. Purity of the total RNA was measured with the NanoDrop 2000 Spectrophotometer. Total RNA was then treated with Turbo DNase (Thermo, #AM2238) with a subsequent RNAClean XP bead cleanup.

### mRNA extraction from yeast cultures

*Saccharomyces cerevisiae* BY4741 (strains: BY4741 MATa pus4::KAN, BY4741 MATa pus1::KAN and BY4741 MATa) were cultured up to log phase at 30°C. The cultures were then divided into two flasks and cultivated at 30°C or 45°C for 1 hour. The cells were harvested via centrifugation at 3,000 rpm for 5 minutes and snap frozen. Total RNA was purified from pelleted cells using a MasterPure Yeast RNA extraction kit (Lucigen, MPY03100), according to manufacturer’s instructions. Total RNA was then DNAse-treated (Ambion, AM2239) at 37°C for 20 minutes with a subsequent clean up using RNeasy MinElute Cleanup Kit (Qiagen, 74204). 70-100 ug of total RNA was subjected to double polyA-selection using Dynabeads Oligo(dT)25 (Invitrogen, 61002) and finally eluted in ice-cold 10 mM Tris pH 7.5.

### Polysome gradient fractionation and rRNA extraction

Yeast pellets from 100 mL cultures were washed with 6 mL of ice-cold Polysome Extraction Buffer (PEB), which contained 20 mM Tris-HCl pH 7.4, 100 mM KCl, 10 mM MgCl_2_, 0.5 mM DTT, 0.1 mg/mL cycloheximide and 100 U/mL RNAse inhibitors (RNaseOUT, Invitrogen, #18080051). Cells were centrifuged for 5 minutes at 3,000 g at 4°C. Washing was repeated by adding 6 mL of ice-cold PEB, followed by centrifugation. Cells were then resuspended in 700 μL of ice-cold PEB, and transferred into pre-chilled 2 mL Eppendorf tubes containing 450 μL of pre-chilled RNAse-free 425-600 μm diameter glass beads (Sigma G8772). Cells were lysed by vortexing at maximum speed for 5 minutes at 4°C, followed by centrifugation also at maximum speed at bench centrifuge for 5 minutes at 4°C. 10% of the supernatant was aliquoted into Trizol for total RNA isolation, and kept at −80°C, which was later used as input. The remaining volume, corresponding approximately to 8 × 10^8^ cells, was subsequently loaded onto the sucrose gradient. Linear sucrose gradients of 10-50% were prepared using the Gradient Station (BioComp). Briefly, SW41 centrifugation tubes (Beckman, Ultra-ClearTM 344059) were filled with Gradient Solution 1 (GS1), which consisted of 20 mM Tris-HCl pH 7.4, 100 mM KCl, 10 mM MgCl_2_, 0.5 mM DTT, 0.1 mg/mL cycloheximide and 10% w/v RNAse-free sucrose. Solutions GS1 and GS2 were prepared with RNase-DNase free UltraPure water and filtered with a 0.22 μM filter. The tube was then filled with 6.3 mL of Gradient Solution 2 (GS2) layered at the bottom of the tube, which consisted of 20 mM Tris-HCl pH 7.4, 100 mM KCl, 10 mM MgCl_2_, 0.5 mM DTT, 0.1 mg/mL cycloheximide and 50% w/v RNAse-free sucrose. The linear gradient was formed using the tilted methodology, with the Gradient Station Maker (Biocomp). Once the gradients were formed, 350 μL of each lysate was carefully loaded on top of the gradients, and tubes were balanced in pairs, placed into pre-chilled SW41Ti buckets and centrifuged at 4°C for 150 minutes at 35,000 rpm. Gradients were then immediately fractionated using the Gradient Station, and 20 × 500 μL fractions were collected in 1.5 mL Eppendorf tubes, while absorbance was monitored at 260 nm continuously. Fractions were combined in the following way: the free rRNA (F1, fractions 1 and 2), the unassembled subunits (F2, fractions 3-6), the lowly-translating monosomes (F3, fractions 7-10) and the highly-translating polysomes (F4, fractions 12-17). The pooled fractions were then concentrated using Amicon-Ultra 100K columns (Millipore), and washed two times with cold PEB. The final volume was brought down to 200 μL, and RNA was extracted using TRIzol reagent. Purity of the RNA was measured with NanoDrop 2000 Spectrophotometer.

### *In vitro* transcription of modified and unmodified RNAs

The synthetic ‘curlcake’ sequences ^29^ used in this study are designed to include all possible 5-mers while minimizing the secondary RNA structure, and consist in 4 *in vitro* transcribed constructs: (i) Curlcake 1, 2244 bp; (ii) Curlcake 2, 2459 bp; (iii) Curlcake 3, 2595 bp, and (iv) Curlcake 4, 2709. The curlcake constructs were *in vitro* transcribed using Ampliscribe™ T7-Flash™ Transcription Kit (Lucigen-ASF3507) with either unmodified rNTPs (UNM), N6-methyladenosine triphosphate (m^6^ATP), 5-methylcytosine triphosphate (m^5^CTP), 5-hydroxymethylcytosine triphosphate (hm^5^CTP) or pseudouridine triphosphate (YTP). All modified NTPs were purchased from TriLink. The sequences included in the short unmodified dataset (UNM-S), which included *B. subtilis* guanine riboswitch, *B. subtilis* lysine riboswitch and *Tetrahymena* ribozyme, were also produced by *in vitro* transcription using Ampliscribe™ T7-Flash™ Transcription Kit (Lucigen-ASF3507). All constructs were 5’ capped using vaccinia capping enzyme (NEB-M2080S) and polyadenylated using *E. coli* Poly(A) Polymerase (NEB-M0276S). Poly(A)-tailed RNAs were purified using RNAClean XP beads, and the addition of poly(A)-tail was confirmed using Agilent 4200 Tapestation. Concentration was determined using Qubit Fluorometric Quantitation. Purity of the IVT product was measured with NanoDrop 2000 Spectrophotometer.

### Direct RNA library preparation and sequencing of *in vitro* transcribed constructs

The RNA libraries for direct RNA Sequencing (SQK-RNA001) were prepared following the ONT Direct RNA Sequencing protocol version DRS_9026_v1_revP_15Dec2016, which corresponds to the flowcell FLO-MIN106. Briefly, 800 ng of Poly(A)-tailed and capped RNA (200 ng per construct) was ligated to ONT RT Adaptor (RTA) using concentrated T4 DNA Ligase (NEB-M0202T), and was reverse transcribed using SuperScript III RT (Thermo Fisher Scientific-18080044). The products were purified using 1.8X Agencourt RNAClean XP beads (Fisher Scientific-NC0068576), washing with 70% freshly prepared ethanol. RNA Adapter (RMX) was ligated onto the RNA:DNA hybrid, and the mix was purified using 1X Agencourt RNAClean XP beads, washing with Wash buffer (WSB) twice. The sample was then eluted in Elution Buffer (ELB) and mixed with RNA running buffer (RRB) prior to loading onto a primed R9.4.1 flowcell, and ran on a MinION sequencer with MinKNOW acquisition software version 1.15.1. The sequencing was performed in independent days and using a different flowcell for each sample (UNM, m^6^A, m^5^C, hm^5^C, Y, UNM-S).

### Direct RNA library preparation and sequencing of yeast total RNAs and mRNAs

Here we performed direct RNA sequencing of two types of *S. cerevisiae* RNA inputs: i) total RNA from *S. cerevisiae*, and ii) polyA-selected RNA from *S. cerevisiae*. Yeast total RNAs were polyadenylated using *E. coli* Poly(A) Polymerase (NEB, M0276S), following the commercial protocol, prior to starting the library prep. Yeast polyA-selected RNA was directly used as input to start the libraries since they already contain poly(A) tail. Four different direct RNA libraries were barcoded according to the recent protocol that we recently published ^92^. Custom RT adaptors (IDT) were annealed using following conditions: custom Oligo A and B (**Table S11**) were mixed in annealing buffer (0.01 M Tris-Cl pH 7.5, 0.05M NaCl) to the final concentration of 1.4 μM each in a total volume of 75 μL. The mixture was incubated at 94°C for 5 minutes and slowly cooled down (−0.1°C/s) to room temperature. RNA library for direct RNA Sequencing (SQK-RNA002) was prepared following the ONT Direct RNA Sequencing protocol version DRS_9080_v2_revI_14Aug2019 with half reaction for each library until the RNA Adapter (RMX) ligation step. Per reaction (half), 250 ng total of yeast RNAs were ligated to pre-annealed custom RT adaptors (IDT) ^92^ using concentrated T4 DNA Ligase (NEB-M0202T), and was reverse transcribed using Maxima H Minus RT (Thermo Scientific, EP0752), without the heat inactivation step. The products were purified using 1.8X Agencourt RNAClean XP beads (Fisher Scientific-NC0068576) and washed with 70% freshly prepared ethanol. 50 ng of reverse transcribed RNA from each reaction was pooled and RMX adapter, composed of sequencing adapters with motor protein, was ligated onto the RNA:DNA hybrid and the mix was purified using 1X Agencourt RNAClean XP beads, washing with Wash Buffer (WSB) twice. The sample was then eluted in Elution Buffer (EB) and mixed with RNA Running Buffer (RRB) prior to loading onto a primed R9.4.1 flowcell, and ran on a MinION sequencer with MinKNOW acquisition software version v.3.5.5.

### NanoCMC-seq

CMC treatment was adapted from Schwartz et al ^4^ with minor changes. Briefly, 20 ug total RNA was incubated in NEBNext^®^ Magnesium RNA Fragmentation Module at 94°C for 1.5 minutes. The fragmented RNA was then incubated with either 0.3 M CMC dissolved in 100 μL TEU buffer (50 mM Tris pH 8.5, 4 mM EDTA, 7 M Urea) or 100 μL TEU buffer (no CMC) for 20 minutes at 37°C. Reaction was stopped with 100 μL of Buffer A (0.3 M NaOAc and 0.1 mM EDTA, pH 5.6), 700 μL absolute ethanol, and 1 μL GlycoBlue (Thermo Scientific, AM9515). RNA in the stop solution was chilled on dry ice for 5 minutes, and then centrifuged at maximum speed for 15 minutes at 4°C. Supernatant was removed and the pellet was washed with 70% ethanol. After air drying for a few minutes, the pellet was dissolved in 100 μL Buffer A and mixed with 300 μL absolute ethanol and 1 μL GlycoBlue. After chilling on dry ice for 5 minutes, the solution was then centrifuged at maximum speed for 15 minutes at 4°C. Supernatant was removed, and the pellet was washed with 70% ethanol. After washing, the pellet was air dried, and resuspended in 40 μL of 50 mM sodium bicarbonate, pH 10.4, and incubated at 37°C for 3 hours. Furthermore, RNA was mixed with 100 μL Buffer A, 700 μl ethanol, and 1 μL Glycoblue overnight at −20°C. The next day, the solution was centrifuged at maximum speed for 15 minutes at 4°C and the pellet was washed with 70% ethanol and dissolved in the appropriate amount of water after air drying. Unprobed and probed RNAs were treated with T4 Polynucleotide Kinase (PNK) (NEB, M0201S) as described above before proceeding with ONT Direct cDNA sequencing.

Before starting the library preparation, 9 μL of 100 μM Reverse-transcription primer (Original ONT VNP: 5’ /5Phos/ACTTGCCTGTCGCTCTATCTTCTTTTTTTTTTTTTTTTTTTTVN 3’) and 9 μL of 100 μM complementary oligo (CompA: 5’ GAAGATAGAGCGACAGGCAAGTA 3’) were mixed with 1 μL 0.2 M Tris pH 7.5 and 1 μL 1 M NaCl. The mix was incubated at 94°C for 1 minute and the temperature was ramped down to 25°C (−0.1°C/s) in order to pre-anneal the oligos. Then, 100 ng polyA-tailed RNA was mixed with 1 μL pre-annealed VNP+CompA, 1 μL 10 mM dNTP mix, 4 μL 5X RT Buffer, 1 μL RNasin^®^ Ribonuclease Inhibitor (Promega, N2511), 1 μL Maxima H Minus RT (Thermo Scientific. EP0742) and nuclease-free water up to 20 μL. The reverse-transcription mix was incubated at 60°C for 60 minutes and inactivated by heating at 85°C for 5 minutes before moving ontoice. Furthermore, RNAse Cocktail (Thermo Scientific, AM2286) was added to the mix in order to digest the RNA and the mix was incubated at 37°C for 10 minutes. Then the reaction was cleaned up using 1.2X AMPure XP Beads (Agencourt, A63881). In order to be able to ligate the sequencing adapters the the first strand, 1 μL 100 μM CompA was again annealed to the 15 μL cDNA in a tube with 2.25 μL 0.1 M Tris pH 7.5, 2.25 μL 0.5 M NaCl and 2 μL nuclease-free water. The mix was incubated at 94°C for 1 minute and the temperature was ramped down to 25 °C (−0.1°C/s) in order to anneal the complementary to the first strand cDNA. Furthermore, 22.5 μL first strand cDNA was mixed with 2.5 μL Native Barcode (EXP-NBD104) and 25 μL Blunt/TA Ligase Mix (NEB, M0367S) and incubated in room temperature for 10 minutes. The reaction was cleaned up using 1X AMPure XP beads and the libraries were pooled into one tube that finally contains 200 fmol library. The pooled library was then ligated to the sequencing adapter (AMII) using Quick T4 DNA Ligase (NEB, M2200S) in room temperature for 10 minutes, followed with 0.65X AMPure XP Bead cleanup using ABB Buffer for washing. The sample was then eluted in Elution Buffer (EB) and mixed with Sequencing Buffer (SQB) and Loading Beads (LB) prior to loading onto a primed R9.4.1 flowcell, and ran on a MinION sequencer with MinKNOW acquisition software version v.3.5.5.

### Analysis of nanoCMC-seq

Reads were base-called with stand-alone Guppy version 3.6.1 with default parameters running in GPU, with built-in demultiplexing tool of Guppy. Unclassified reads were then demultiplexed further using Porechop with --barcode_threshold 50 option (https://github.com/rrwick/Porechop). Then all the merged classified reads were mapped to cytosolic and mitochondrial ribosomal RNA sequences in S. cerevisiae using minimap2 default. Furthermore, a custom script was used to extract RT-drop signatures and the RT-drop scores were plotted using ggplot2. All scripts used to process nanoCMC-seq data with RT-Drop information have been made available in GitHub (https://github.com/novoalab/yeast_RNA_Mod). Notably, due to the 5’ end truncation of the nanopore sequencing reads by ~13 nt, RT-drop positions were shifted by 13 nt to accurately determine the exact RT-drop positions. To identify significant RT drops in a given transcript, we first computed RT-drop scores at each site, which took the difference in the coverage at a given position (0) relative to the previous position (−1). We then computed the difference (delta RT drop-off score) in RT-drop scores between CMC-probed and unprobed conditions. Lastly, we normalized the delta RT drop-off score at each position by the median RT drop-off per transcript, leading to final CMC-Scores, which can be compared across transcripts. Positions with CMC-Score greater than 25 were considered significant, i.e. to contain a pseudouridine. We should note that the nanoCMC-seq signal-to-noise ratio is dependent on the coverage of the individual transcript.

### Demultiplexing direct RNA sequencing

Demultiplexing of the barcoded direct RNA sequencing libraries was performed using DeePlexiCon with default parameters ^92^. Reads with demultiplexing confidence scores greater than 0.95 were kept for downstream analyses. We used a lower score in the case of polysomal fractions and mRNA runs (0.8), due to the low read coverage of some fractions and/or genes. We should note that the dataset was also analyzed using 0.95 threshold, and results and conclusions of the analysis did not change, compared to those obtained using 0.80 threshold.

### Base-calling direct RNA sequencing

Reads were base-called with stand-alone Albacore versions 2.1.7 and 2.3.4 with the --disable_filtering parameter, and stand-alone Guppy versions 2.3.1 and 3.0.3 with default parameters running in CPU. In-house scripts were used for computing the number of unique and common base-called reads between the different approaches, as well as to compare the tendency of each base-caller regarding read lengths and qualities. Both Albacore and Guppy are available to ONT customers via their community site (https://community.nanoporetech.com/). Differences between the base-called features using distinct base-callers were determined using Kruskal-Wallis test with Bonferroni correction for pairwise comparisons, whereas differences between unmodified and modified sites were assessed using Mann-Whitney-Wilcoxon test.

### Mapping algorithms and parameters

Reads were mapped using either *Minimap2* ^44^ or *GraphMap* ^45^. *Minimap2* version 2.14 was run with two different parameter settings: (i) minimap2 -ax map-ont, which is the recommended setting for direct RNA sequencing mapping, and thus we refer to as ‘default’, and (ii) minimap2 -ax map-ont -k 5, which we refer to as ‘sensitive’. *GraphMap* version 0.5.2 was also run with two different parameter settings, for comparison, (i) graphmap align, using ‘default’ parameters, and (ii) graphmap align -- rebuild-index -v 1 --double-index --mapq -1 -x sensitive -z 1 -K fastq --min-read-len 0 -A 7 -k 5, which is expected to increase the tolerance to errors that may occur under the presence of RNA modifications, and thus we refer to as ‘sensitive’. Yeast total RNA runs were mapped to ribosomal RNAs and non-coding RNA transcripts using graphmap with default settings. Yeast poly(A)-selected runs were mapped to the yeast genome (SacCer3) using minimap2 with -ax splice -k14 -uf parameters. The scripts can be found in the GitHub repository https://github.com/novoalab/yeast_RNA_Mod. Sequencing, base-calling and mapping statistics for all yeast sequencing runs (total RNA and polyA-selected RNA) can be found in **Tables S12** and **13**.

### Analysis of base-called features in curlcakes

Sam files were transformed into bam files using Samtools version 1.9 ^93^, and were then sorted and indexed in order to visualize the data using the Integrative Genomics Viewer (IGV) version 2.4.16 ^94^. Base-called features were extracted with *EpiNano* version 1.1 (https://github.com/enovoa/EpiNano). Principal Component Analysis (PCA) was used to reduce the dimensionality of the base-calling error data to visually inspect for base-calling differences, using as input the base-called features (mismatch frequency, deletion frequency and per-base quality) from all 5 positions of each k-mer. Only k-mers that contained a given modification once in the 5-mer were included in the analysis. All scripts used to analyze *in vitro* transcribed sequences using different base-calling algorithms and mappers, as well as to generate the Figures related to their analysis are available in https://github.com/novoalab/Best_Practices_dRNAseq_analysis.

### Analysis of base-called features in yeast RNAs

Sam files were transformed into bam files using Samtools version 1.9 ^93^, then sorted and indexed in order to visualize the data using the Integrative Genomics Viewer (IGV) version 2.4.16 ^94^. Base-called features were extracted using *EpiNano* version 1.1 with minor modifications, which consisted in including in the output csv file the directionality of mismatched bases (C_frequency, G_frequency, A_frequency, U_frequency). The modified *EpiNano* script can be found at https://github.com/novoalab/yeast_RNA_Mod. Scripts for the analysis and visualization of base-called features are also included in the same GitHub repository.

### Visualization per-read current intensities using *Nanopolish*

*Nanopolish* eventalign output was processed to extract the current intensity values corresponding to the 15-mer regions centered in the modified sites, for the following sites: (i) 6 Y rRNA sites for which knockout data was available (25s:2133, 25s:2129, 25s:2826, 25s:2880, 25s:2264, 18s:1187), for all 4 sequencing datasets (wild type, snR3-KO, snR34-KO, snR36-KO); (ii) 4 Nm sites for which knockout data was available (25s:817, 25s:908, 25s:1133, 25s:1888), for all 4 sequencing datasets (wild type, snR60-KO, snR61-KO, snR62-KO); (iii) 7 Y snRNA/snoRNA sites which were identified as heat-sensitive, for which there was a minimum of 100 reads of coverage. Reads with empty values in the 15-mer region in the *Nanopolish* eventalign output were omitted from the analysis.

### Analysis of current intensity, dwell time and trace

In this work, we used two different softwares to extract current intensity: Nanopolish ^95^ and Tombo ^49^. Nanpolish was used to extract the aligned current intensity values per read and position, using the option *--scale-events*. Mean current intensity per-position was computed by summing the current intensities of all reads aligned to the same position, divided by the total number of reads mapping at a given position. All scripts used to process *Nanopolish* event align output, including scripts to display mean current intensity values along transcripts have been made available in GitHub (https://github.com/novoalab/nanoRMS).

Signal intensity, dwell time and trace were retrieved using get_features.py script, which is available as part of *nanoRMS*. This program internally uses: minimap2 (read alignment), Tombo (calculation of signal intensity and dwell time) and ont-fast5-api (retrieval of trace). Trace represents the probability that a given signal intensity chunk may be originating from each of the 4 canonical bases (A, C, G and T/U), and it is reported relative to the reference base. For example, in a T reference position that is incorrectly reported as C (common base-calling error observed for Y sites), the trace value will be reported for the reference base (T in this case). Then, the final read alignment and all the features are stored into sorted BAM files. All scripts necessary to retrieve and store per-read, per-position features and plot/calculate results are available within the *nanoRMS* GitHub repository (https://github.com/novoalab/nanoRMS).

### *De novo* prediction of pseudouridine modifications on yeast mitochondrial rRNAs

To systematically identify Y sites *de novo* based on the Y base-calling signatures, we first extracted the mismatch frequency and per-base mismatch frequency (C_freq, A_freq, U_freq, G_freq) from both unmodified (U) and modified (Y) sites from cytosolic ribosomal RNAs, from three biological replicates. As expected, C mismatch frequency (C_freq) and global mismatch frequency (mis_freq) showed clearly distinct distributions when comparing unmodified and Y-modified sites (**Figure 4A**). We then determined the optimal cut-points for these two features using the *cutpointr* package in R with oc_youden_kernel method, which applies Kernel smoothing and maximizes the Youden-Indexing. This approach predicted C_freq=0.137 and mis_freq= 0.587 as optimal cut-offs. For the mitochondrial ribosomal RNA, we filtered the uridine sites based on the selected features and assigned those that are replicable in three biological replicates as “candidate” pseudouridine sites.

### *De novo* prediction of pseudouridine modifications in yeast mRNAs and non-coding RNAs

Due to the lower stoichiometry of modification of noncoding RNAs (snRNA and snoRNAs) and mRNAs, we focused on analysis of the *de novo* detection of Y sites whose pseudouridylation levels would be changing between two conditions, either by comparing normal and stress (heat-shock) conditions, or by comparing the base-calling ‘error’ patterns of wild type strains and Pus1 or Pus4-deficient strains. Only sites which passed the coverage filter (n>30 reads) in both biological replicates from both conditions were considered in the analysis (**Table S6**). Sites with minimal mismatch frequency difference of 0.1 between the two conditions in both replicates that met the identified Y signature (C_freq=0.137 and mis_freq= 0.587) were considered as true Y sites that were either heat-sensitive, Pus1-dependent, or Pus4-dependent, respectively. The individual candidate Y mRNA and ncRNA sites identified using nanopore sequencing, as well as the previously reported Y mRNA and ncRNA sites (using CMC probing coupled to Illumina sequencing) can be found in **Tables S7-S10**.

### Prediction of RNA modification stoichiometry using *nanoRMS*

Per-position features from individual reads were stored in BAM files using pysam (https://github.com/pysam-developers/pysam) and stored them either in Numpy arrays (https://numpy.org/) or Pandas DataFrames (https://pandas.pydata.org/) using the script get_features.py, which is available as part of *nanoRMS*. Models were trained with combinations of features with diverse ranges of sequence contexts surrounding the modified sites (k=1-15). Features used to predict stoichiometry included: (i) current intensity (SI), (ii) dwell time in the centre of the pore (at position 0, DT/DT0), (iii) dwell time at helicase centre (shifted by 10 positions, DT10) and (iv) base probability (trace, TR). Estimation of modification frequency was performed using unsupervised (GMM, KMEANS, IsolationForest, OneClassSVM) and supervised (KNN, RandomForest) machine learning methods implemented in sklearn (https://sklearn.org/). Plots were built using matplotlib and seaborn (https://seaborn.pydata.org/).

Trained models were first benchmarked with unmodified (KO) and modified (WT) reads from rRNA mutants dataset, to identify which machine learning methods and which combination of features discriminated between modified and unmodified reads. Then, we tested how the diverse models would perform at diverse stoichiometries of modification. To this end, we simulated samples with varying levels of modification: 0%, 20%, 40%, 60%, 80% and 100% (using mixes of KO and WT reads) and estimated the modification level in those simulated samples by comparing them to KO (**Figure S5C**).

*NanoRMS* performed best when trained with signal intensity (SI) + trace (TR) as features, and when using KNN supervised models or KMEANS unsupervised models, both for Y and Nm-modified sites. Predictions by each clustering algorithm, and for each individual rRNA modified site, are shown in **Table S4**. For mRNA and ncRNA analysis, only sites with more than 30 reads of coverage in all conditions and replicates were included for predicting RNA modification stoichiometry. Prediction of RNA modification stoichiometry in mRNAs and non-coding RNAs was performed using signal intensity + trace as features, and k-means as classification algorithm. Stoichiometry changes were reported as the difference in predicted stoichiometry between the two conditions. All code and examples to predict RNA modification stoichiometry are available as part of the *nanoRMS* GitHub repository (https://github.com/novoalab/nanoRMS).

## DATA AVAILABILITY

For *in vitro* transcribed datasets, FAST5 files used in this work were already publicly available (UNM and m^6^A: PRJNA521324), or have been made publicly available in SRA (m^5^C:PRJNA563591; hm^5^C: PRJNA548268; Y:PRJNA511582, UNM-S: PRJNA575545). Base-called and demultiplexed FASTQ from all yeast RNA direct RNA sequencing data runs have been made publicly available in GEO, under the accession number GSE148603, including processed *EpiNano* outputs and *Nanopolish* outputs. FAST5 files for all yeast RNA direct RNA sequencing are available in ENA under accession PRJEB37798 and PRJEB41495. A detailed description of the datasets used and sequenced in this work, with their corresponding GEO and ENA/SRA IDs can be found in **Table S14**. BAM files with extracted features for the rRNA mapped reads in WT and snoRNA-depleted strains are available through the *nanoRMS* GitHub repository (https://github.com/novoalab/nanoRMS).

## CODE AVAILABILITY

All scripts and code used in this work have been made available in GitHub: https://github.com/novoalab/Best_Practices_dRNAseq_analysis (analysis of *in vitro* curlcake datasets), https://github.com/novoalab/yeast_RNA_Mod (analysis of *in vivo* datasets) and https://github.com/novoalab/nanoRMS (prediction of RNA modifications and estimation of RNA modification stoichiometries).

## ACKNOWLEDGEMENTS

We thank all the members of the Novoa lab for their valuable insights and discussion. We thank Vivek Malhotra for sharing the Pus1 and Pus4 knockout strains. OB is supported by a UNSW International PhD fellowship. MCL is supported by an FPI Severo-Ochoa fellowship by the Spanish Ministry of Economy, Industry and Competitiveness (MEIC). IM and SC are supported by “la Caixa” INPhINIT PhD fellowships (LCF/BQ/DI18/11660028 and LCF/BQ/DI19/11730036, respectively). This project has received funding from the European Union’s Horizon 2020 research and innovation programme under the Marie Skodowska-Curie grant agreement No. 713673. This work was supported by the Australian Research Council (DP180103571 to EMN) and the Spanish Ministry of Economy, Industry and Competitiveness (MEIC) (PGC2018-098152-A-100 to EMN). We acknowledge the support of the MEIC to the EMBL partnership, Centro de Excelencia Severo Ochoa and CERCA Programme/Generalitat de Catalunya.

## AUTHOR CONTRIBUTIONS

OB and MCL performed the majority of wet lab experiments, including RNA extraction and nanopore library preparation. OB and LPP performed bioinformatic analysis of the data, together with JMR and EMN. OB conceived and performed nanoCMC-Seq experiments. MCL produced the *in vitro* transcribed sequences with modifications and their corresponding nanopore libraries. OB produced the *in vitro* transcribed sequences with different pseudouridine stoichiometry and performed their corresponding nanopore library. LPP benchmarked and wrote the *nanoRMS* code, together with OB and EMN. JMR performed bioinformatic analyses on *in vitro* transcribed constructs and compared base-calling and mapping algorithms. IM built polysome gradients and helped with their corresponding nanopore libraries. SC and IM prepared and sequenced the 2’-O-methylation mutant strains. HGSV and RM cultured the *S. cerevisiae* strains under different stress conditions. HL contributed with code for the analysis of current intensity values. ASC cultured all snoRNA-depleted yeast mutant strains and extracted their total RNA. EMN conceived the project. EMN supervised the work, with the assistance of SS and JSM. MCL, OB and EMN built the figures. OB, MCL and EMN wrote the paper, with contributions from all authors.

## REFERENCES

1. Dominissini, D. et al. Topology of the human and mouse m6A RNA methylomes revealed by m6A-seq. Nature 485, 201–206 (2012).

2. Meyer, K. D. et al. Comprehensive Analysis of mRNA Methylation Reveals Enrichment in 3′ UTRs and near Stop Codons. Cell vol. 149 1635–1646 (2012).

3. Carlile, T. M. et al. Pseudouridine profiling reveals regulated mRNA pseudouridylation in yeast and human cells. Nature 515, 143–146 (2014).

4. Schwartz, S. et al. Transcriptome-wide mapping reveals widespread dynamic-regulated pseudouridylation of ncRNA and mRNA. Cell 159, 148–162 (2014).

5. Lovejoy, A. F., Riordan, D. P. & Brown, P. O. Transcriptome-wide mapping of pseudouridines: pseudouridine synthases modify specific mRNAs in S. cerevisiae. PLoS One 9, e110799 (2014).

6. Li, X. et al. Chemical pulldown reveals dynamic pseudouridylation of the mammalian transcriptome. Nat. Chem. Biol. 11, 592–597 (2015).

7. Hussain, S., Aleksic, J., Blanco, S., Dietmann, S. & Frye, M. Characterizing 5-methylcytosine in the mammalian epitranscriptome. Genome Biol. 14, 215 (2013).

8. Huang, T., Chen, W., Liu, J., Gu, N. & Zhang, R. Genome-wide identification of mRNA 5-methylcytosine in mammals. Nat. Struct. Mol. Biol. 26, 380–388 (2019).

9. Delatte, B. et al. RNA biochemistry. Transcriptome-wide distribution and function of RNA hydroxymethylcytosine. Science 351, 282–285 (2016).

10. Safra, M. et al. The m1A landscape on cytosolic and mitochondrial mRNA at single-base resolution. Nature 551, 251–255 (2017).

11. Li, X. et al. Base-Resolution Mapping Reveals Distinct mA Methylome in Nuclear- and Mitochondrial-Encoded Transcripts. Mol. Cell 68, 993–1005.e9 (2017).

12. Marchand, V. et al. AlkAniline-Seq: Profiling of m7G and m3C RNA Modifications at Single Nucleotide Resolution. Angew. Chem. Int. Ed. 57, 16785–16790 (2018).

13. Arango, D. et al. Acetylation of Cytidine in mRNA Promotes Translation Efficiency. Cell 175, 1872–1886.e24 (2018).

14. Sas-Chen, A. et al. Dynamic RNA acetylation revealed by quantitative cross-evolutionary mapping. Nature (2020) doi:10.1038/s41586-020-2418-2.

15. Zhang, L.-S. et al. Transcriptome-wide Mapping of Internal N7-Methylguanosine Methylome in Mammalian mRNA. Mol. Cell 74, 1304–1316.e8 (2019).

16. Pandolfini, L. et al. METTL1 Promotes let-7 MicroRNA Processing via m7G Methylation. Mol. Cell 74, 1278–1290.e9 (2019).

17. Delaunay, S. & Frye, M. RNA modifications regulating cell fate in cancer. Nat. Cell Biol. 21, 552–559 (2019).

18. Haussmann, I. U. et al. m6A potentiates Sxl alternative pre-mRNA splicing for robust Drosophila sex determination. Nature vol. 540 301–304 (2016).

19. Vu, L. P. et al. The N6-methyladenosine (m6A)-forming enzyme METTL3 controls myeloid differentiation of normal hematopoietic and leukemia cells. Nature Medicine vol. 23 1369–1376 (2017).

20. Novoa, E. M., Mason, C. E. & Mattick, J. S. Charting the unknown epitranscriptome. Nat. Rev. Mol. Cell Biol. 18, 339–340 (2017).

21. Anreiter, I., Mir, Q., Simpson, J. T., Janga, S. C. & Soller, M. New Twists in Detecting mRNA Modification Dynamics. Trends Biotechnol. 0, (2020).

22. Li, X., Xiong, X. & Yi, C. Epitranscriptome sequencing technologies: decoding RNA modifications. Nat. Methods 14, 23–31 (2016).

23. Motorin, Y. & Helm, M. Methods for RNA Modification Mapping Using Deep Sequencing: Established and New Emerging Technologies. Genes 10, (2019).

24. Grozhik, A. V. et al. Antibody cross-reactivity accounts for widespread appearance of m 1 A in 5’UTRs. Nat. Commun. 10, 1–13 (2019).

25. Lahens, N. F. et al. IVT-seq reveals extreme bias in RNA sequencing. Genome Biol. 15, R86 (2014).

26. Garalde, D. R. et al. Highly parallel direct RNA sequencing on an array of nanopores. Nat. Methods 15, 201–206 (2018).

27. Jonkhout, N. et al. The RNA modification landscape in human disease. RNA 23, 1754–1769 (2017).

28. Leger, A., Amaral, P. P., Pandolfini, L. & Capitanchik, C. RNA modifications detection by comparative Nanopore direct RNA sequencing. BioRxiv (2019).

29. Liu, H. et al. Accurate detection of m6A RNA modifications in native RNA sequences. Nat. Commun. 10, 4079 (2019).

30. Parker, M. T. et al. Nanopore direct RNA sequencing maps the complexity of Arabidopsis mRNA processing and m6A modification. eLife vol. 9 (2020).

31. Price, A. M. et al. Direct RNA sequencing reveals m6A modifications on adenovirus RNA are necessary for efficient splicing. bioRxiv 865485 (2019) doi:10.1101/865485.

32. Wongsurawat, T. et al. Decoding the Epitranscriptional Landscape from Native RNA Sequences. bioRxiv 487819 (2018) doi:10.1101/487819.

33. Lorenz, D. A., Sathe, S., Einstein, J. M. & Yeo, G. W. Direct RNA sequencing enables m6A detection in endogenous transcript isoforms at base-specific resolution. RNA 26, 19–28 (2020).

34. Jack, K. et al. rRNA pseudouridylation defects affect ribosomal ligand binding and translational fidelity from yeast to human cells. Mol. Cell 44, 660–666 (2011).

35. Yoon, A. et al. Impaired control of IRES-mediated translation in X-linked dyskeratosis congenita. Science 312, 902–906 (2006).

36. Bellodi, C. et al. Loss of Function of the Tumor Suppressor DKC1 Perturbs p27 Translation Control and Contributes to Pituitary Tumorigenesis. Cancer Research vol. 70 6026–6035 (2010).

37. Wu, G., Xiao, M., Yang, C. & Yu, Y.-T. U2 snRNA is inducibly pseudouridylated at novel sites by Pus7p and snR81 RNP. EMBO J. 30, 79–89 (2011).

38. Taoka, M., Nobe, Y., Hori, M. & Takeuchi, A. A mass spectrometry-based method for comprehensive quantitative determination of post-transcriptional RNA modifications: the complete chemical structure of …. Nucleic acids (2015).

39. Basu, A. et al. Requirement of rRNA methylation for 80S ribosome assembly on a cohort of cellular internal ribosome entry sites. Mol. Cell. Biol. 31, 4482–4499 (2011).

40. Marcel, V. et al. p53 acts as a safeguard of translational control by regulating fibrillarin and rRNA methylation in cancer. Cancer Cell 24, 318–330 (2013).

41. Belin, S. et al. Dysregulation of Ribosome Biogenesis and Translational Capacity Is Associated with Tumor Progression of Human Breast Cancer Cells. PLoS ONE vol. 4 e7147 (2009).

42. Buchhaupt, M. et al. Partial methylation at Am100 in 18S rRNA of baker’s yeast reveals ribosome heterogeneity on the level of eukaryotic rRNA modification. PLoS One 9, e89640 (2014).

43. Chen, H. et al. METTL5, an 18S rRNA-specific m6A methyltransferase, modulates expression of stress response genes. bioRxiv 2020.04.27.064162 (2020) doi:10.1101/2020.04.27.064162.

44. Li, H. Minimap2: pairwise alignment for nucleotide sequences. Bioinformatics vol. 34 3094–3100 (2018).

45. Sović, I. et al. Fast and sensitive mapping of nanopore sequencing reads with GraphMap. Nat. Commun. 7, 11307 (2016).

46. Liu, Q. et al. Detection of DNA base modifications by deep recurrent neural network on Oxford Nanopore sequencing data. Nat. Commun. 10, 2449 (2019).

47. McIntyre, A. B. R. et al. Single-molecule sequencing detection of N6-methyladenine in microbial reference materials. Nat Commun 10: 579. (2019).

48. De Coster, W., Stovner, E. B. & Strazisar, M. Methplotlib: analysis of modified nucleotides from nanopore sequencing. Bioinformatics 36, 3236–3238 (2020).

49. Stoiber, M. et al. De novo Identification of DNA Modifications Enabled by Genome-Guided Nanopore Signal Processing. bioRxiv 094672 (2017) doi:10.1101/094672.

50. Taoka, M. et al. The complete chemical structure of Saccharomyces cerevisiae rRNA: partial pseudouridylation of U2345 in 25S rRNA by snoRNA snR9. Nucleic Acids Res. 44, 8951–8961 (2016).

51. Pintard, L., Bujnicki, J. M., Lapeyre, B. & Bonnerot, C. MRM2 encodes a novel yeast mitochondrial 21S rRNA methyltransferase. EMBO J. 21, 1139–1147 (2002).

52. Sharma, S. & Lafontaine, D. L. J. ‘View From A Bridge’: A New Perspective on Eukaryotic rRNA Base Modification. Trends Biochem. Sci. 40, 560–575 (2015).

53. Taoka, M. et al. Landscape of the complete RNA chemical modifications in the human 80S ribosome. Nucleic Acids Res. 46, 9289–9298 (2018).

54. Fischer, N. et al. Structure of the E. coli ribosome””EF-Tu complex at< 3 Å resolution by C s-corrected cryo-EM. Nature 520, 567–570 (2015).

55. Sergeeva, O. V., Bogdanov, A. A. & Sergiev, P. V. What do we know about ribosomal RNA methylation in Escherichia coli? Biochimie 117, 110–118 (2015).

56. Sloan, K. E. et al. Tuning the ribosome: The influence of rRNA modification on eukaryotic ribosome biogenesis and function. RNA Biol. 14, 1138–1152 (2017).

57. Hebras, J., Krogh, N., Marty, V., Nielsen, H. & Cavaillé, J. Developmental changes of rRNA ribose methylations in the mouse. RNA Biol. 1–15 (2019).

58. Higa-Nakamine, S. et al. Loss of ribosomal RNA modification causes developmental defects in zebrafish. Nucleic Acids Res. 40, 391–398 (2012).

59. Sahoo, T. et al. Prader-Willi phenotype caused by paternal deficiency for the HBII-85 C/D box small nucleolar RNA cluster. Nat. Genet. 40, 719–721 (2008).

60. Heiss, N. S. et al. X-linked dyskeratosis congenita is caused by mutations in a highly conserved gene with putative nucleolar functions. Nat. Genet. 19, 32–38 (1998).

61. Knight, S. W. et al. X-Linked Dyskeratosis Congenita Is Predominantly Caused by Missense Mutations in the DKC1 Gene. The American Journal of Human Genetics vol. 65 50–58 (1999).

62. Liao, J. et al. Small nucleolar RNA signatures as biomarkers for non-small-cell lung cancer. Mol. Cancer 9, 198 (2010).

63. Mei, Y.-P. et al. Small nucleolar RNA 42 acts as an oncogene in lung tumorigenesis. Oncogene 31, 2794–2804 (2012).

64. Bortolin-Cavaille, M.-L., -L. Bortolin-Cavaille, M. & Cavaille, J. The SNORD115 (H/MBII-52) and SNORD116 (H/MBII-85) gene clusters at the imprinted Prader-Willi locus generate canonical box C/D snoRNAs. Nucleic Acids Research vol. 40 6800–6807 (2012).

65. Erales, J. et al. Evidence for rRNA 2’-O-methylation plasticity: Control of intrinsic translational capabilities of human ribosomes. Proc. Natl. Acad. Sci. U. S. A. 114, 12934–12939 (2017).

66. Krogh, N. et al. Profiling of 2′-O-Me in human rRNA reveals a subset of fractionally modified positions and provides evidence for ribosome heterogeneity. Nucleic Acids Research vol. 44 7884–7895 (2016).

67. Birkedal, U. et al. Profiling of ribose methylations in RNA by high-throughput sequencing. Angew. Chem. Int. Ed Engl. 54, 451–455 (2015).

68. Natchiar, S. K., Myasnikov, A. G., Kratzat, H., Hazemann, I. & Klaholz, B. P. Visualization of chemical modifications in the human 80S ribosome structure. Nature 551, 472–477 (2017).

69. van der Feltz, C., DeHaven, A. C. & Hoskins, A. A. Stress-induced Pseudouridylation Alters the Structural Equilibrium of Yeast U2 snRNA Stem II. J. Mol. Biol. 430, 524–536 (2018).

70. Frye, M., Harada, B. T., Behm, M. & He, C. RNA modifications modulate gene expression during development. Science 361, 1346–1349 (2018).

71. Roundtree, I. A., Evans, M. E., Pan, T. & He, C. Dynamic RNA Modifications in Gene Expression Regulation. Cell 169, 1187–1200 (2017).

72. Li, S. & Mason, C. E. The pivotal regulatory landscape of RNA modifications. Annu. Rev. Genomics Hum. Genet. 15, 127–150 (2014).

73. Wang, X. et al. LARP7-Mediated U6 snRNA Modification Ensures Splicing Fidelity and Spermatogenesis in Mice. Mol. Cell 77, 999–1013.e6 (2020).

74. Louloupi, A., Ntini, E., Conrad, T. & Ørom, U. A. V. Transient N-6-Methyladenosine Transcriptome Sequencing Reveals a Regulatory Role of m6A in Splicing Efficiency. Cell Reports vol. 23 3429–3437 (2018).

75. Zhou, K. I. et al. Regulation of Co-transcriptional Pre-mRNA Splicing by m6A through the Low-Complexity Protein hnRNPG. Mol. Cell 76, 70–81.e9 (2019).

76. Lee, Y., Choe, J., Park, O. H. & Kim, Y. K. Molecular Mechanisms Driving mRNA Degradation by m6A Modification. Trends Genet. 36, 177–188 (2020).

77. Guo, M., Liu, X., Zheng, X., Huang, Y. & Chen, X. m6A RNA Modification Determines Cell Fate by Regulating mRNA Degradation. Cellular Reprogramming vol. 19 225–231 (2017).

78. Geula, S. et al. Stem cells. m6A mRNA methylation facilitates resolution of naïve pluripotency toward differentiation. Science 347, 1002–1006 (2015).

79. Weng, H. et al. METTL14 Inhibits Hematopoietic Stem/Progenitor Differentiation and Promotes Leukemogenesis via mRNA m6A Modification. Cell Stem Cell 22, 191–205.e9 (2018).

80. Lence, T. et al. m6A modulates neuronal functions and sex determination in Drosophila. Nature 540, 242–247 (2016).

81. Kan, L. et al. The m6A pathway facilitates sex determination in Drosophila. Nat. Commun. 8, 15737 (2017).

82. Schaefer, M., Kapoor, U. & Jantsch, M. F. Understanding RNA modifications: the promises and technological bottlenecks of the ‘epitranscriptome’. Open Biology vol. 7 170077 (2017).

83. Depledge, D. P. et al. Direct RNA sequencing on nanopore arrays redefines the transcriptional complexity of a viral pathogen. Nat. Commun. 10, 754 (2019).

84. Roach, N. P. et al. The full-length transcriptome of C. elegans using direct RNA sequencing. Genome Res. 30, 299–312 (2020).

85. Workman, R. E. et al. Nanopore native RNA sequencing of a human poly(A) transcriptome. doi:10.1101/459529.

86. Kim, D. et al. The Architecture of SARS-CoV-2 Transcriptome. Cell 181, 914–921.e10 (2020).

87. Pratanwanich, P. N. et al. Detection of differential RNA modifications from direct RNA sequencing of human cell lines. bioRxiv 2020.06.18.160010 (2020) doi:10.1101/2020.06.18.160010.

88. Bokar, J. A., Shambaugh, M. E., Polayes, D., Matera, A. G. & Rottman, F. M. Purification and cDNA cloning of the AdoMet-binding subunit of the human mRNA (N6-adenosine)-methyltransferase. RNA 3, 1233–1247 (1997).

89. Torchet, C. et al. The complete set of H/ACA snoRNAs that guide rRNA pseudouridylations in Saccharomyces cerevisiae. RNA 11, 928–938 (2005).

90. Hamma, T. & Ferré-D’Amaré, A. R. Pseudouridine synthases. Chem. Biol. 13, 1125–1135 (2006).

91. Parker, S. et al. A resource for functional profiling of noncoding RNA in the yeast Saccharomyces cerevisiae. RNA 23, 1166–1171 (2017).

92. Smith, M. A. et al. Barcoding and demultiplexing Oxford Nanopore native RNA sequencing reads with deep residual learning. bioRxiv 864322 (2019) doi:10.1101/864322.

93. Li, H. et al. The Sequence Alignment/Map format and SAMtools. Bioinformatics vol. 25 2078–2079 (2009).

94. Robinson, J. T. et al. Integrative genomics viewer. Nat. Biotechnol. 29, 24–26 (2011).

95. Loman, N. J., Quick, J. & Simpson, J. T. A complete bacterial genome assembled de novo using only nanopore sequencing data. Nat. Methods 12, 733–735 (2015).

