## Supplementary Figures for "Quantitative profiling of native RNA modifications and their dynamics using nanopore sequencing"

**Figure S1. Bench-marking of base-calling and mapping algorithms enables dissection of RNA modification base-calling 'error' signatures and reveals their sequence context-dependence. (A)** IGV snapshots of unmodified (UNM), $m^6A$-modified ($m^6A$), $m^5C$-modified ($m^5C$), $hm^5C$-modified ($hm^5C$) or Y-modified (Y) *in vitro* transcribed sequence Curlcake 1, base-called using either Albacore 2.1.7 (AL 2.1.7) or Guppy 3.0.3 (GU 3.0.3), and then mapped using minimap2 or GraphMap in 'sensitive' mode. Nucleotides with mismatch frequencies greater than 0.1 have been colored. **(B)** Mean sequence identity of different combinations of base-calling and mapping algorithms, for each of the 6 *in vitro* transcribed datasets analyzed. **(C)** Comparison of read lengths and per-read mean quality scores in different *in vitro* transcribed datasets (UNM, $m^6A$, Y, $m^5C$, $hm^5C$ and UNM-S) when base-called using different algorithms (AL 2.1.7, AL 2.3.4, GU 2.3.1 or GU 3.0.3). Results show that read lengths do not largely vary across base-callers. By contrast, per-read quality strongly varies depending on the choice of base-calling algorithm. Box, first to last quartiles; whiskers, 1.5x interquartile range; center line, median; points, outliers. **(D)** Barplots of mean per-read quality show that per-read qualities are slightly decreased in all modified datasets, relative to unmodified ones, with this difference being most evident in GU 3.0.3 base-called data. **(E)** Boxplots of mean per-base quality of reads base-called with GU 3.0.3 show that per-base qualities are decreased in all modified datasets, relative to unmodified ones. Box, first to last quartiles; whiskers, 1.5x interquartile range; center line, median; points, outliers. **(F)** Ternary plots depicting the mismatch distribution of the unmodified (left) and modified (right) positions colored by log coverage, in 5 different datasets: unmodified (all left panels), $m^6A$-modified ($m^6A$), Y-modified (Y), $m^5C$-modified ($m^5C$), $hm^5C$-modified ($hm^5C$). Only modified nucleotides, and their relative unmodified counterparts in the UNM dataset, are shown. Each dot represents a different nucleotide in the reference. **(G)** Logo representations of the mismatch signatures generated by $m^5C$ and $hm^5C$. Results show that the signatures are different depending on the modification, however, these also vary depending on the 5-mer sequence (reported on the left).

**Figure S1** (legend in previous page)

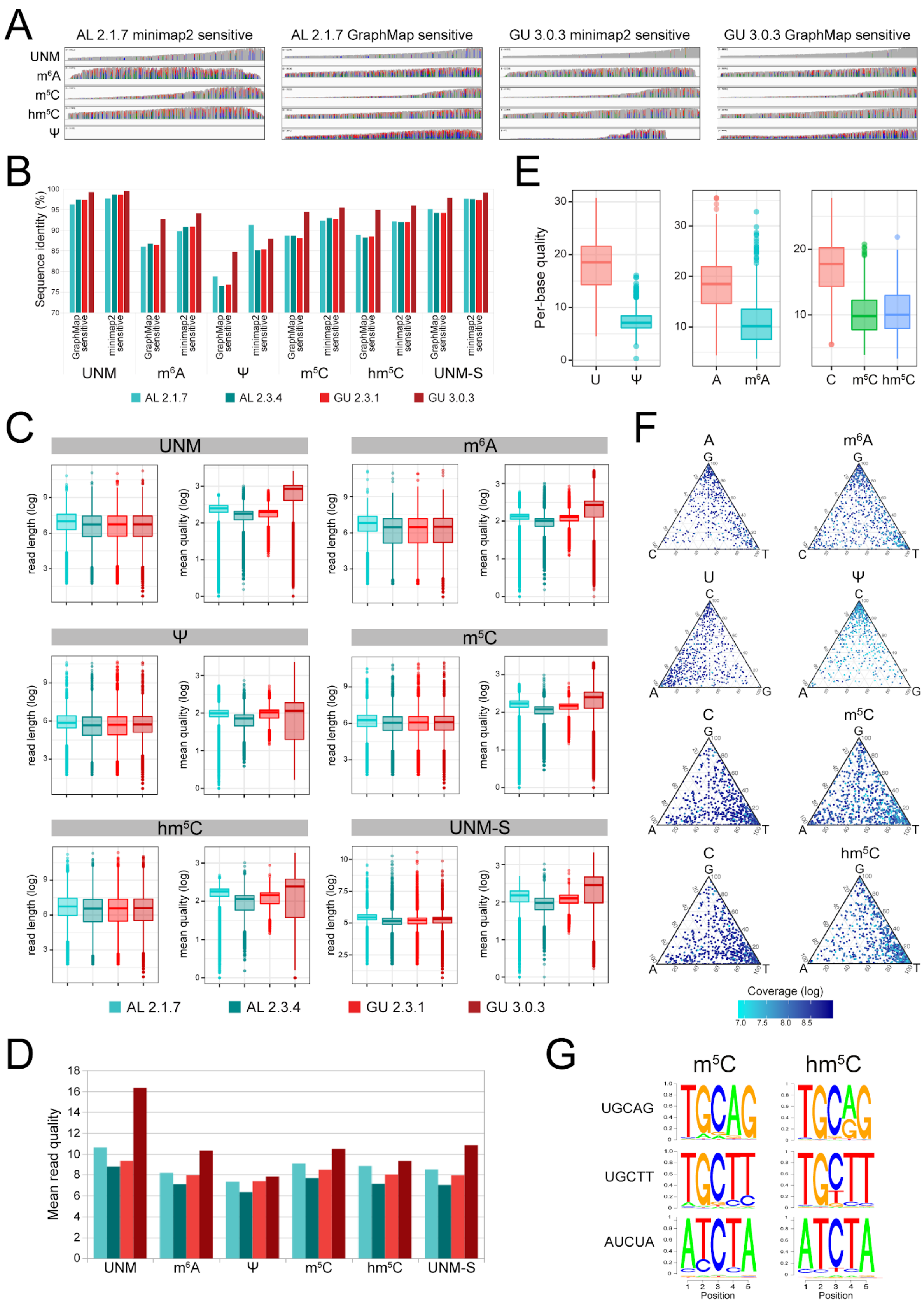

**Figure S2. Known yeast ribosomal RNA modifications show distinct base-calling 'error' signatures. (A)** IGV snapshots centered in distinct yeast ribosomal RNA modifications in 4 different yeast strains (wild type, snR3-KO, snR34-KO, snR36-KO, in descending order). Known rRNA modification sites are indicated below each snapshot. Nucleotides with mismatch frequencies greater than 0.15 have been colored. **(B)** Dotplots of base-calling errors (deletion frequency, insertion frequency, mismatch frequency, and per-base quality) observed in modified 5-mers, centered in the modified position. Each dot corresponds to a different 5-mer. The total number of 5-mers included in the analysis varies depending on the abundance of each rRNA modification type in yeast rRNAs: Y (n=46), Am (n=14), Cm (n=10), Gm (n=15) and Um (n=9). 5-mers that contain more than one modification in the 5-mer region were excluded from the analysis. Box, first to last quartiles; whiskers, 1.5x interquartile range; center line, median; points, individual data points.

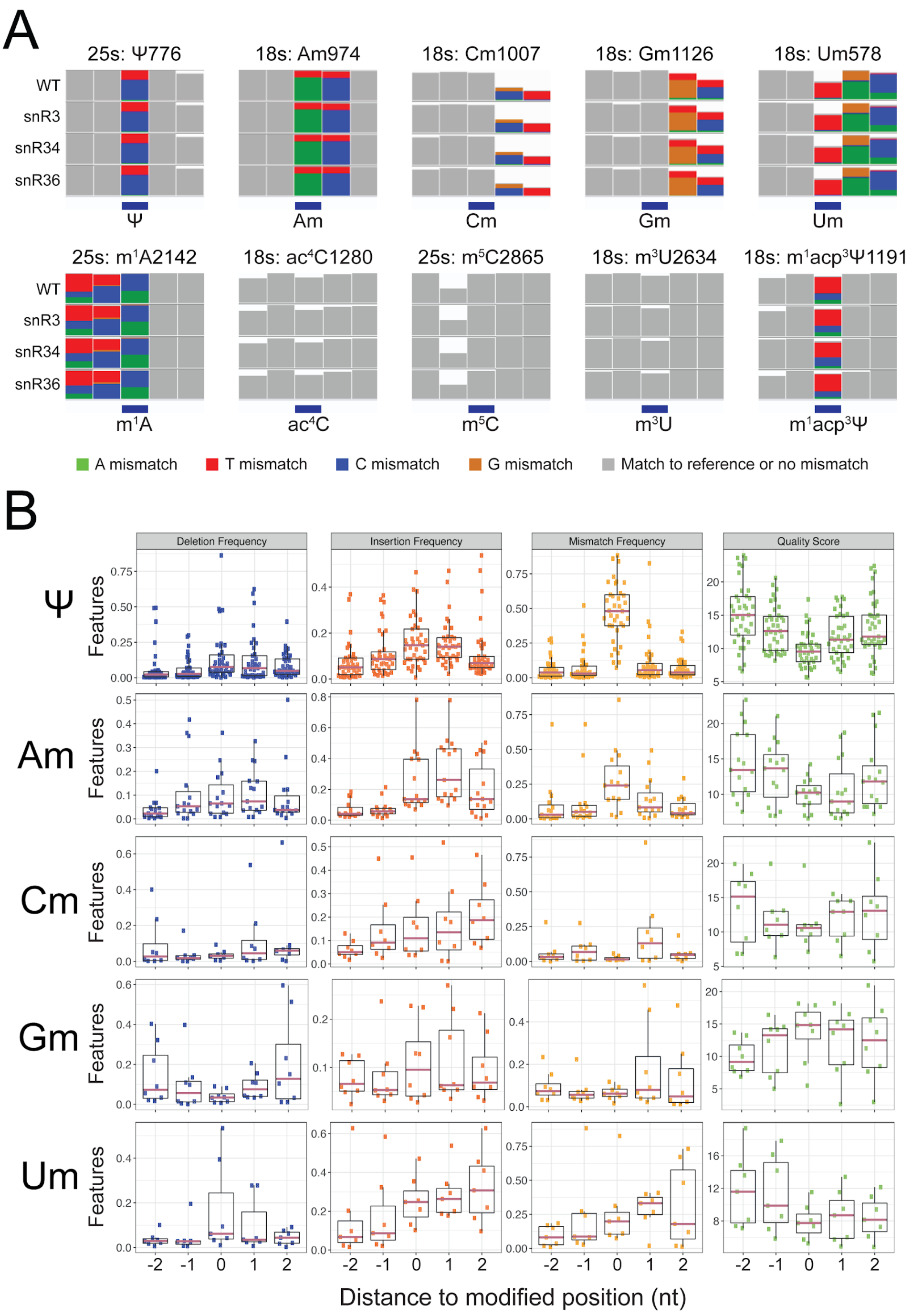

**Figure S3. Base-calling signature of 2'-O-methylations often alter the neighboring positions, whereas Y modifications mainly affect the modified site. (A)** IGV snapshots centered on known yeast rRNA modified sites: Y-modified sites are shown in the upper panels, whereas 2'-O-methylated sites are shown in the bottom panels. Nucleotides with mismatch frequencies greater than 0.15 have been colored. **(B)** Comparison of base-calling 'errors' (mismatch, deletion and insertion frequency) observed in snoRNA-depleted strains (snR60, top panels; snR61, middle panels, snR62, bottom panels) relative to wild type, with snoRNA target sites indicated in red, neighboring sites indicated in blue and non-target sites in gray.

**Figure S3** (legend in previous page)

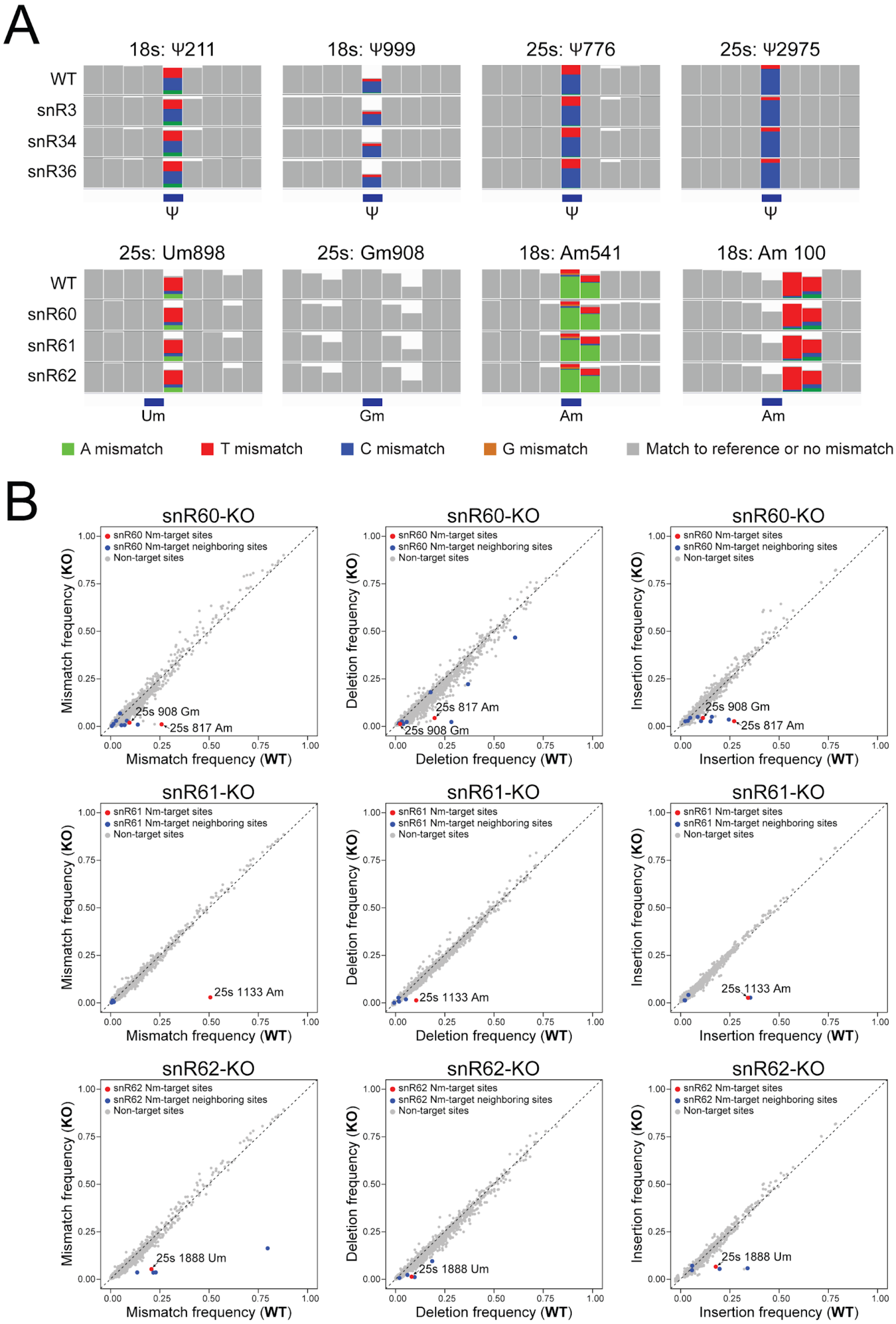

**Figure S4. Pseudouridylations and 2'-O-methylations can be detected in the form of altered current intensities. (A)** Distributions of per-read current intensity at known Y-modified, 2'-O-methylated and negative control sites. Y and 2'-O-methylated positions were altered upon deletion of specific snoRNAs relative to wild type, whereas no shift was observed in control sites. **(B)** Absolute differences in current intensity along the 25s rRNA and 18s rRNAs upon depletion of snR34 and snR36, respectively, relative to the wild type strain. Red vertical lines indicate the KO pseudouridylation positions. **(C)** Comparison of mean current intensity changes for Y and 2'-O-methyl knockout sites across each of the snoRNA knockout strains. The dotted vertical line indicates the modified position. **(D)** Per-read analysis of current intensities centered at 3 different Y modified sites targeted by the snoRNAs depleted in each knockout strain (25s:Y2129, 25s:Y2264 and 18s:Y1187). In each panel, the per-read current intensities centered in the modified site are shown, both for the wild type (purple) and knockout strain (red: snR3; green: snR34; cyan: snR36). As a control, the same analysis was performed at a control site (25s:Y986), using reads from wild type (purple) and snR34 knockout strain (green) showing no differences between the read populations. Each line indicates a single read. **(E)** Principal Component Analysis of the current intensity values of the 15-mer regions was performed, and the corresponding scatterplots of the two first principal components (PC1 and PC2) are depicted for 4 different Y and Nm sites. Each dot corresponds to a different read, and is colored according to the strain.

**Figure S4** (legend in previous page)

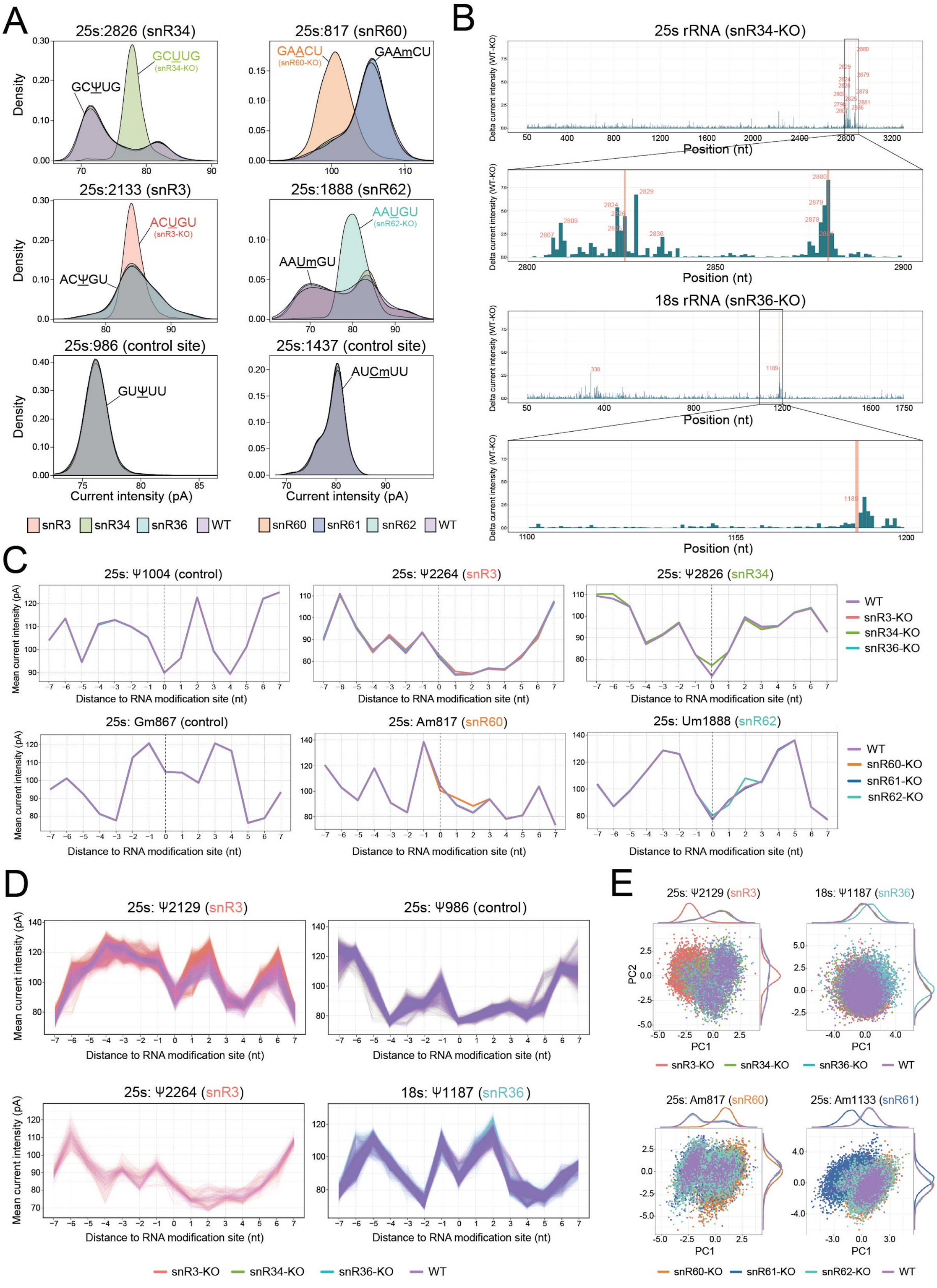

**Figure S5. Systematic benchmarking of resquiggling softwares, machine learning algorithms and distinct feature sets for the prediction of RNA modification stoichiometry from individual RNA reads. (A)** Comparative analysis of read resquiggling using *Nanopolish* and *Tombo,* depicting the relative proportion of resquiggled reads for each algorithm at each individual site. The Y-modified sites and 2'-O-methylated sites were analyzed independently, as they come from independent flowcells. *Tombo* shows uniform proportion of resquiggled reads along the same transcript, whereas *Nanopolish* shows variable proportion of resquiggled reads depending on the site. **(B)** Comparative analysis of read resquiggling using *Nanopolish* and *Tombo*, depicting the relative proportion of resquiggled reads from KO strains for a given position (relative to WT), using as input 1000 reads for each strain, and for each algorithm. If there is no difference in resquiggling depending on the presence or absence of modification, the expected proportion of KO:WT reads is 1. **(C)** Line chart of expected (X-axis) and observed (Y-axis) modification frequency for Y-modified and 2'-O-methylated positions. The absolute modification frequency difference was estimated between two samples: KO (no modified reads) and WT (simulating varying levels of modification frequency: 0.0, 0.2, 0.4, 0.6, 0.8 and 1.0). Modification stoichiometry was calculated using four different machine learning methods: two supervised (k-nearest neighbour (KNN) and random forest (RF)) and two unsupervised (K-means and gaussian mixture model (GMM). Five sets of distinct feature combinations have been tested for each algorithm and RNA modification type: current intensity (blue), trace (orange), dwell time (green), the combination of current intensity and trace (red) and the combination of current intensity and dwell time (purple).

**Figure S5** (legend in previous page)

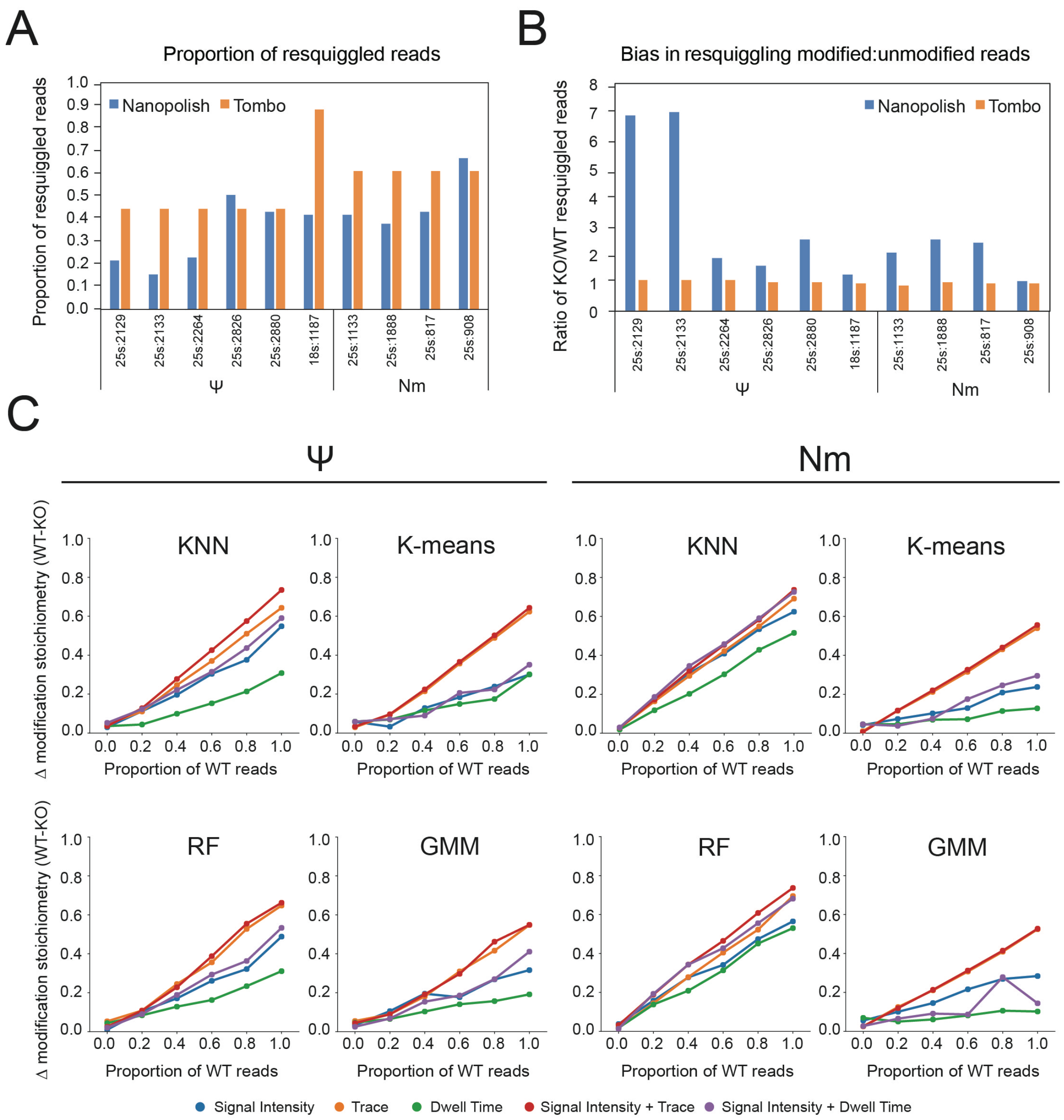

**Figure S6. Density plots of the per-read current intensity, trace and dwell time features in selected Y and 2'-O-methylated rRNA sites. (A,B)** Per-read distributions of current intensity (SI), trace (TR) and dwell time (DT) between respective mutants and wild type at Y-modified **(A)**, and 2'-O-methylated sites **(B)**. Control sites, which are not affected by any of the knockouts, have been included in the analysis as negative controls. The distributions are plotted for the positions of interest (0) and two neighbouring positions: downstream (-1) and upstream (+1). The density distribution of each feature has been colored depending on the strain. X-axis scale depends on the feature type: SI is reported as median absolute deviation normalised signal intensity as reported by *Tombo*; TR is reported as reference-base probability (0-1 scaled), and DT is reported as log2 (observed/expected), where expected is dwell time mean value per base calculated for every read.

**Figure S6** (legend in previous page)

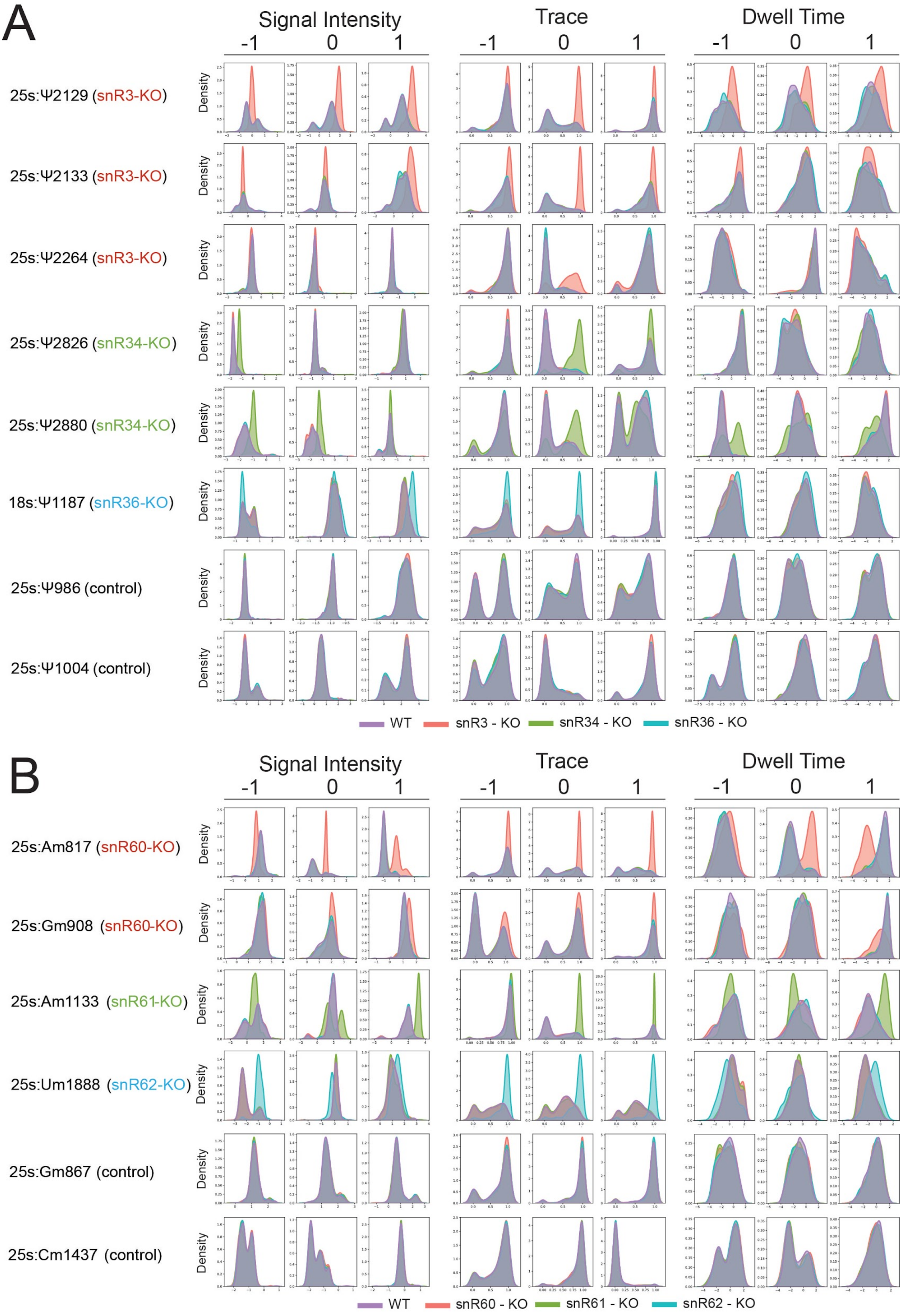

**Figure S7.** ***De novo*** **prediction of Y modifications reveals a novel Pus4-dependent modification (15s:Y854) in yeast mitochondrial rRNAs, and captures previously reported Pus4-dependent mRNA modifications. (A)** NanoCMC-seq scores along the 21s mitochondrial LSU rRNA (upper panel) and the 25s cytosolic LSU rRNA (bottom panel). Dashed lines indicate the CMC-score threshold used for determining the positive sites. All the positions with a significant CMC Score (>25) correspond to known Y rRNA modification sites (blue). **(B)** Comparison of mismatch frequency for each base in Pus4 knockout strains, relative to wild type, in positions mapped to yeast genome and rRNA, in two independent biological replicates. **(C)** IGV snapshots of wild type (rep1 and rep2) and Pus4 knockout (rep1 and rep2) yeast strains with zoomed subsets depicting the site-specific loss of mismatch at known target positions. Nucleotides with mismatch frequencies greater than 0.15 have been colored. **(D)** Stress scores in previously reported heat-responsive sn/snoRNA pseudouridylated sites (as defined by Schwartz et al 2014). Stress scores are calculated by taking the difference between mismatch frequency in stress (heat-shock, cold-shock, and oxidative) and normal conditions. (**E)** Stress scores in sn/snoRNA Y sites that were not previously reported as heat-responsive. Our analysis identifies some of these sites as responsive to heat-stress. **(F)** Polysome profiles of ribosomal-bound RNA fractions isolated from untreated and stressed $H_2O_2$-treated yeast cells. **(G)** Comparison of mismatch frequency for untreated vs $H_2O_2$-treated input RNA (upper panel) and untreated vs $H_2O_2$-treated ribosome-bound RNA (lower panel). **(H)** Comparison of mismatch frequencies of ribosomal RNAs for different fractions (F1: Free, F2: Subunit, F3: Monosome, F4: Polysome). Each dot represents a base in the rRNA, with significantly altered Y sites reproducible across biological replicates highlighted in red. The remaining Y sites are shown in black and the rest of the sites in gray. All rRNA bases from cytosolic rRNAs were included in the analysis and plots.

**Figure S7** (legend in previous page)

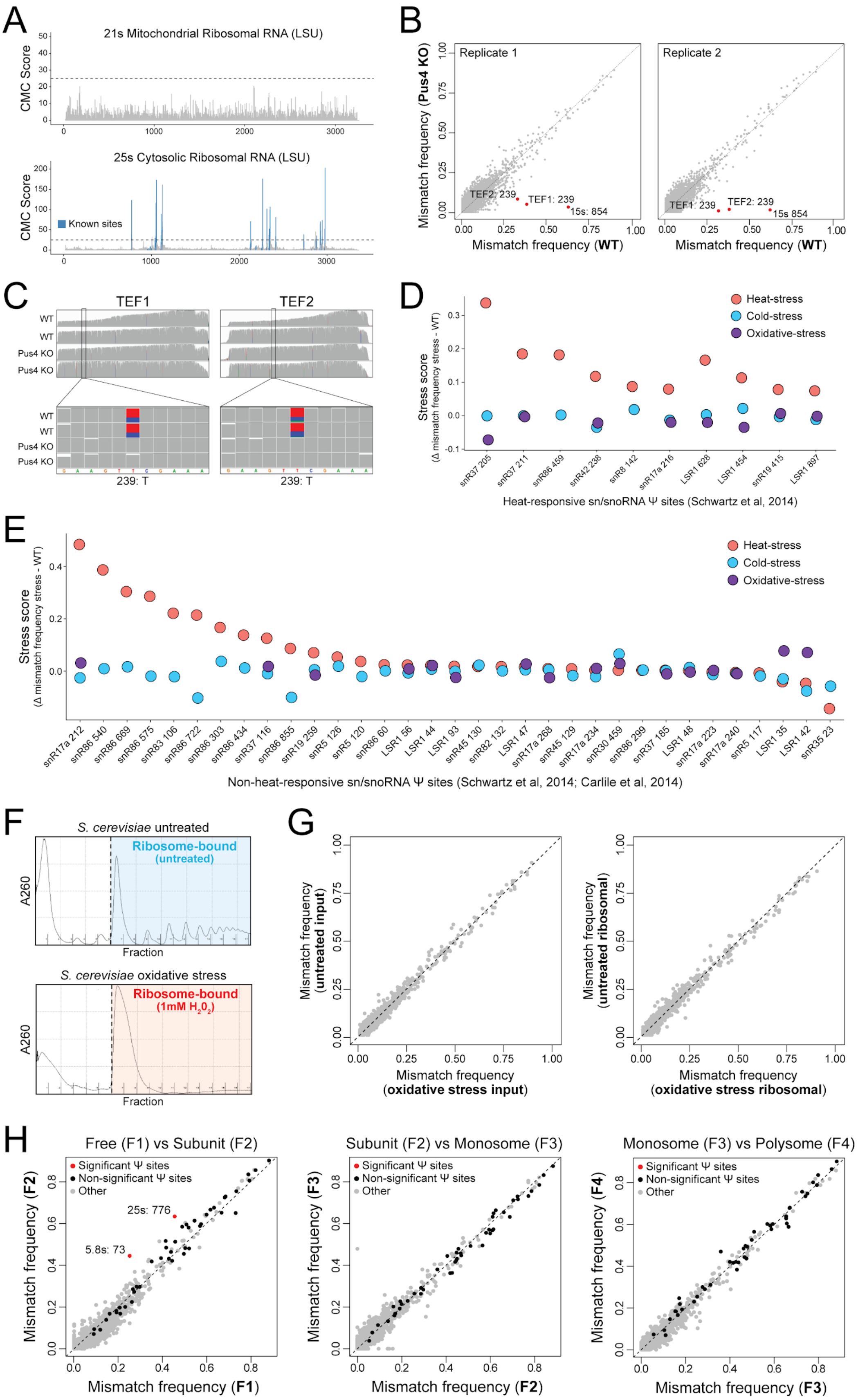

**Figure S8. Analysis of features in previously reported and novel mRNA Y sites. (A,B)** Per-read distributions of current intensity (SI), trace (TR) and dwell time (DT) in Pus1 KO and wild type *S. cerevisiae* strains (A) and Pus4 KO and wild type strains (B), for 4 different Y sites (reported and novel). The distributions are shown for the features observed at the modified site (0) as well as at the two neighbouring positions: downstream (-1) and upstream (+1), both in predicted and not predicted Y sites. The density of each feature has been colored depending on the sample. X-axis scale depends on the feature type: SI is reported as median absolute deviation normalised signal intensity as reported by *Tombo*; TR is reported as reference-base probability (0-1 scaled), and DT is reported as log2 (observed/expected), where expected is dwell time mean value per base. **(B)** Comparison of per-read distributions of current intensity (SI), trace (TR) and dwell time (DT) in normal (30°C) and heat (45°C) conditions, at 4 different Y sites (reported and novel). **(C)** Venn diagrams illustrate the overlap of predicted Y sites in mRNAs and ncRNAs between two studies (Schwartz et al, 2014 and Carlile et al, 2014). **(D)** Current intensity density plots showing altered current intensity distribution in positions 25s:Y2826 (left panel) and 25s:Y2880 (right panel), in the wild type strain (which corresponds to Y-centered k-mers) and snR34 knockout strain (which corresponds to U-centered k-mers). Current intensity distribution of the equivalent 5-mers with C in the middle position are shown in blue.

**Figure S8** (legend in previous page)

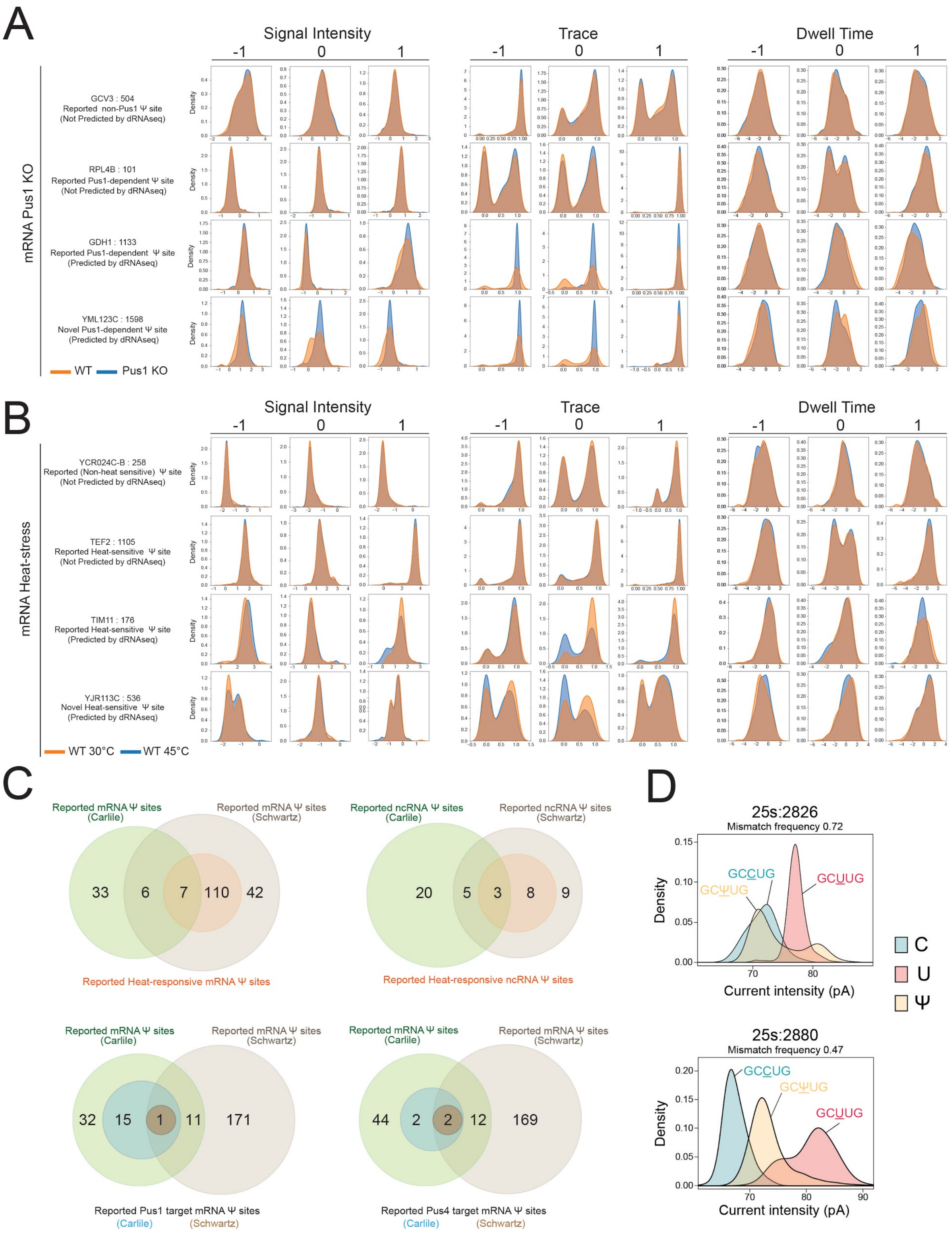
